# Proteomic and functional comparison between human induced and embryonic stem cells

**DOI:** 10.1101/2021.10.20.464767

**Authors:** Alejandro J. Brenes, Eva Griesser, Linda V. Sinclair, Lindsay Davidson, Alan R. Prescott, Francois Singh, Elizabeth K.J. Hogg, Carmen Espejo-Serrano, Hao Jiang, Harunori Yoshikawa, Melpomeni Platani, Jason Swedlow, Greg M. Findlay, Doreen A. Cantrell, Angus I. Lamond

## Abstract

Human induced pluripotent stem cells (hiPSCs) have great potential to be used as alternatives to embryonic stem cells (hESCs) in regenerative medicine and disease modelling, thereby avoiding many of the ethical issues arising from the use of embryo-derived cells. However, despite clear similarities between the two cell types, it is likely they are not identical. In this study, we characterise the proteomes of multiple hiPSC and hESC lines derived from independent donors. We find that while hESCs and hiPSCs express a near identical set of proteins, they show consistent quantitative differences in the expression levels of a wide subset of proteins.

hiPSCs have increased total protein content, while maintaining a comparable cell cycle profile to hESCs. The proteomic data show hiPSCs have significantly increased abundance of vital cytoplasmic and mitochondrial proteins required to sustain high growth rates, including nutrient transporters and metabolic proteins, which correlated with phenotypic differences between hiPSCs and hESCs. Thus, higher levels of glutamine transporters correlated with increased glutamine uptake, while higher levels of proteins involved in lipid synthesis correlated with increased lipid droplet formation. Some of the biggest metabolic changes were seen in proteins involved in mitochondrial metabolism, with corresponding enhanced mitochondrial potential, shown experimentally using high-resolution respirometry. hiPSCs also produced higher levels of secreted proteins, including ECM components and growth factors, some with known tumorigenic properties, as well as proteins involved in the inhibition of the immune system. Our data indicate that reprogramming of human fibroblasts to iPSCs effectively restores protein expression in cell nuclei to a state comparable to hESCs, but does not similarly restore the profile of cytoplasmic and mitochondrial proteins, with consequences for cell phenotypes affecting growth and metabolism. The data improve understanding of the molecular differences between induced and embryonic stem cells, with implications for potential risks and benefits for their use in future disease modelling and therapeutic applications.

## Introduction

Human embryonic stem cells (hESC) are derived from the inner cell mass of a pre-implantation embryo^1^. They show prolonged undifferentiated potential, as well as the ability to differentiate into the three main embryonic germ layers^2^, making them excellent models for studying disease mechanisms, development and differentiation. However, their use remains restricted by regulations, based in part upon ethical considerations^3^.

Over a decade ago, methods allowing the induction of pluripotent stem cells from fibroblast cultures, in both human and mice, were developed^4,5^. These reports showed that by exogenously expressing a small set of key transcription factors (Oct4, Sox2, c-Myc and Klf4), a somatic cell could be reprogrammed back into a pluripotent state, characterised by their capacity for self-renewal and ability to differentiate into the three main germ layers. These human induced pluripotent stem cells (hiPSCs) show many key features of their physiological embryonic stem cell (hESC) counterparts, while avoiding many of the ethical issues regarding the use of stem cells derived from embryos.

Since the discovery of reprogramming methods, hiPSC lines have attracted great interest, particularly for their potential use as alternatives to hESCs in regenerative medicine^6^ and disease modelling, including studies on monogenic disorders^7,8^ and some late onset diseases^9^. However, to understand the value of using hiPSCs in regenerative therapy, drug development and/or studies of disease mechanisms, it is important to establish how similar hiPSCs are to hESCs at the molecular and functional levels. To address this, multiple studies have compared hiPSCs and hESCs, using a variety of assays, including methylation analysis^10^, transcriptomics^11,12^ and even quantitative proteomics^13,14^. It should be noted, however, that many of these earlier studies were performed at a time when reprogramming protocols were less robust^15^ and when the depth of proteome coverage and quantitative information that could be obtained was lower than today.

In this study, we have addressed the similarity of hiPSCs to hESCs by performing a detailed proteomic analysis, comparing a set of four hiPSC lines derived from human primary skin fibroblasts^16^ of independent, healthy donors, with four independent hESC lines. The data highlight that while both types of stem cell lines have very similar global protein abundance profiles, they also show some specific and significant quantitative differences in protein expression. In particular, the reprogramed iPSC lines consistently display higher total protein levels, predominantly affecting cytoplasmic proteins required to sustain higher growth, along with mitochondrial changes, and an excess of secreted proteins, with impact upon cell phenotypes.

## Results

### hESCs and hiPSCs display quantitative differences in protein abundances

For this study, we compared multiple hESC and hiPSC lines, all derived from different donors and cultured using identical growth conditions. First, the expression levels of the main pluripotency markers were verified in each of the lines, with no differences detected between the respective hESC and hiPSC cell types (Fig. 1a). From these data, representative sets of four hiPSCs and four hESCs lines were selected for detailed proteomic analysis using mass spectrometry. The proteomes were characterised using tandem mass tags (TMT)^17^, within a single 10-plex (Table S1) and using MS3-based synchronous precursor selection^18^ (SPS). To further optimise quantification accuracy, each sample was allocated to a specific isobaric tag to minimise cross-population reporter ion interference (Fig. 1b), as previously described^19^. In total 8,491 protein groups (henceforth referred to as ‘proteins’), were detected at 1% FDR, with >99% overlap between the proteins detected from both the hESC and hiPSC lines (Fig. 1c). However, it is important to highlight that TMT is not the best method to use when looking for proteins that are specific to one condition or population, as protein detected in one channel are frequently seen across all other channels as well^19^.

**Figure 1-.**
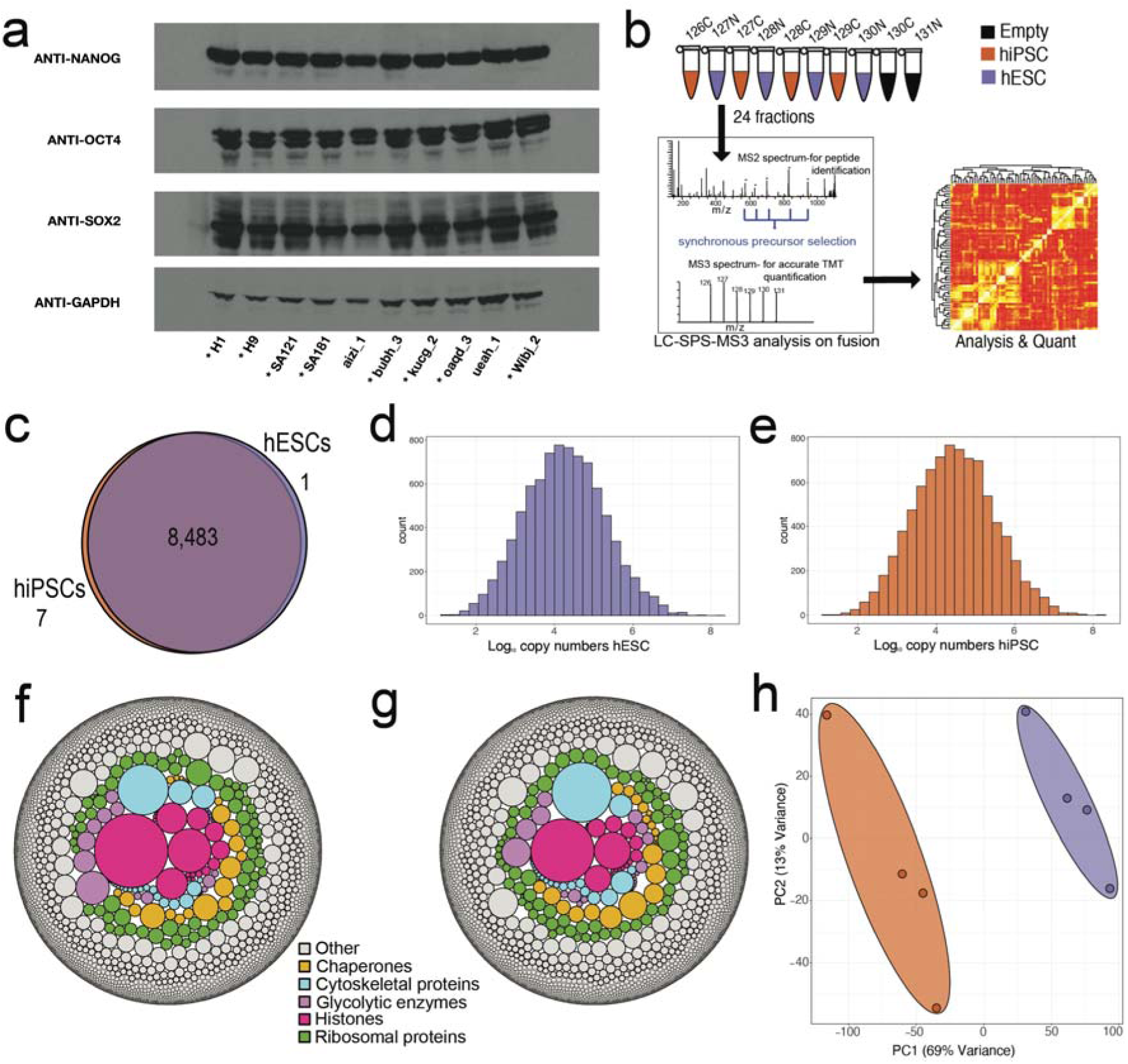
Proteomic overview: **(a)** Western blots showing the expression of the pluripotency factors NANOG, OCT4 and SOX2 across all hESC and hiSPC lines. The 8 lines showed with * were used within the proteomic analysis **(b)** Diagram showing the SPS-MS3 TMT proteomic workflow used for the experiment. **(c)** Venn diagram showing the overlap of proteins identified within the hiPSC and hESC populations. **(d)** Average copy number histogram for the hESCs. (N=4) **(e)** Average copy number histogram for the hiPSCs (N=4). **(f)** Bubble plot showing proteins coloured by specific categories where the size is represented by the average hESC estimated protein copy numbers. **(g)** Bubble plot showing proteins coloured by specific categories where the size is represented by the average hiPSC estimated protein copy numbers. **(h)** PCA plot based on the log_10_ copy numbers for all 8 stem cell lines. hESCs are shown in purple and hiPSCs in orange.

To provide a quantitative comparison of the respective proteomes, we focussed on analysing the 7,878 proteins that were detected with at least 2 unique and razor peptides (see methods). After confirming that there were no differences in the abundance levels of histones between the two cell types (Fig. S1), protein copy numbers were estimated via the “proteomic ruler”^20^ (see methods). The copy number data (Supplemental Table 1) highlighted that both the hESC (Fig. 1d) and hiPSC (Fig. 1e), proteomes display a similar dynamic range, with estimated protein copy numbers extending from a median of less than 100 copies, to over 100 million copies per cell. Furthermore, the composition of the respective proteomes is highly similar. Both cell types display high expression levels of ribosomal proteins, protein chaperones and glycolytic enzymes (Fig. 1f&g), consistent with rapid proliferation and dependence on glycolysis for energy generation^21^. It is only when the quantitative data are examined in more detail that differences between the cell types become apparent (Fig. 1h). A principal component analysis (PCA), based on the protein copy numbers, revealed a clear separation between the two stem cell populations within the main component of variation, which accounted for 69% of variance. The PCA suggested that the independent hiPSC lines were clearly different to the hESC lines, and vice versa.

### Standard normalisation methods mask changes in total protein content in hiPSCs compared to hESCs

A previous proteomic study reported that there were virtually no protein level differences between hESCs and hiPSCs^13^. However, in that study the intensity data were median normalised. We therefore decided to compare two different normalisation methods: i.e., the previously used median normalisation method and the “proteomic ruler”^20^. The median normalisation produces concentration-like results and is frequently used to normalise proteomic data. With this approach, our data also show no major differences in protein abundances between the hESC and hiPSC lines (Fig. 2a; Supplemental Table 2), i.e. ~94% of all proteins displayed no significant changes in abundance (FC>1.5-fold; q-value < 0.001), similar to the previously reported conclusion^13^. However, median (or total intensity) normalisation methods lack the capacity to detect changes in absolute abundance, cell size or protein content. By artificially forcing all medians to be almost identical, such changes are invisible.

**Figure 2-.**
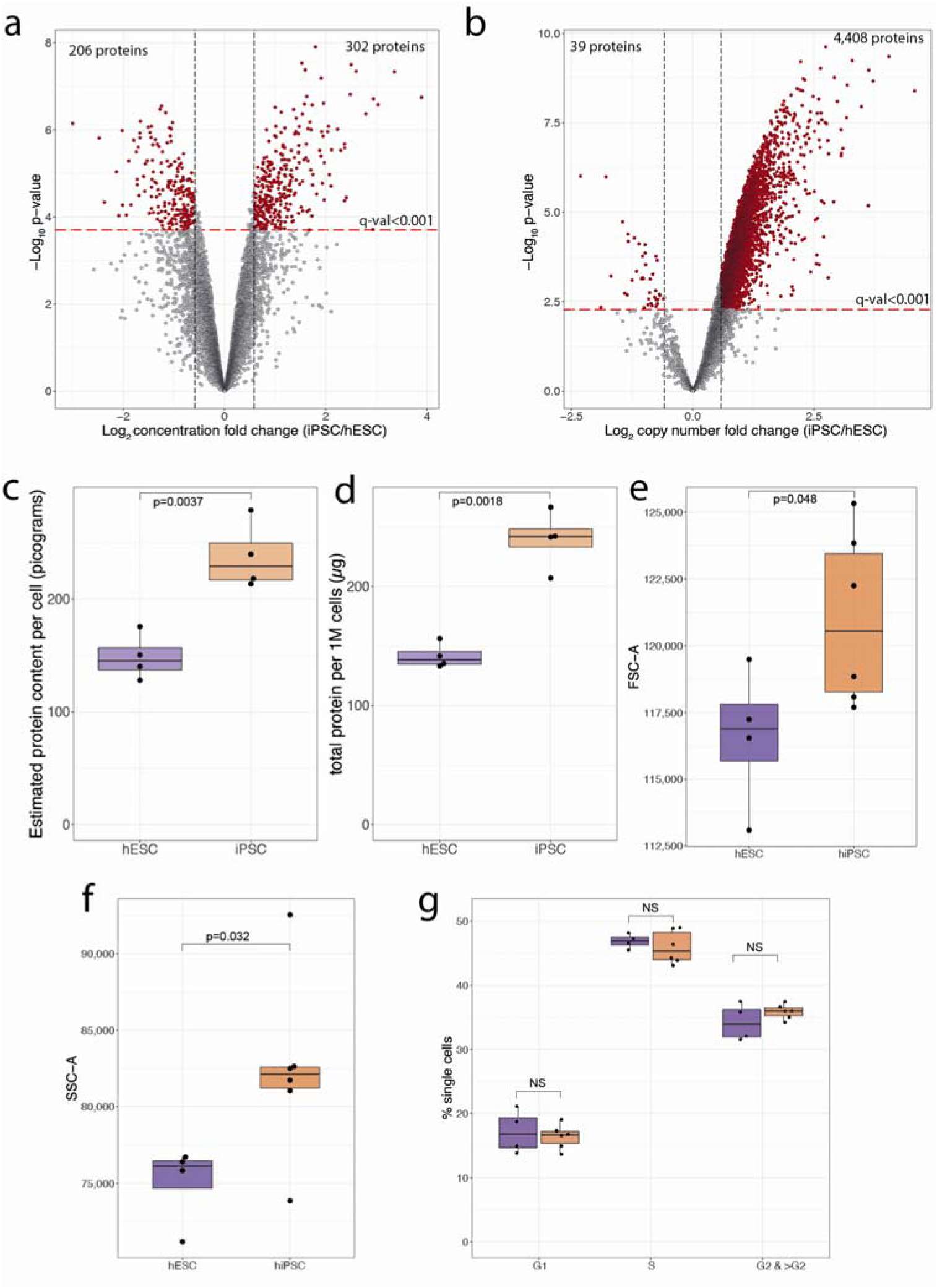
Normalisation and protein content: **(a)** Concentration based volcano plot showing the −log_10_ p-value and the log_2_ fold change comparing hiPSCs (N=4) to hESCs (N=4). Elements shaded in red are considered significantly changed. All dots above the red line have a q-value lower than 0.001 **(b)** Copy number-based volcano plot showing the −log_10_ p-value and the log_2_ fold change comparing hESCs (N=4) to hiSPC (N=4). Elements shaded in red are considered significantly changed. All dots above the red line have a q-value lower than 0.001 **(c)** Box plot showing the MS based estimated protein content for hESCs (N=4) and hiPSC(N=4). **(d)** Box plot showing the protein amount per million cells derived from the EZQ Protein Quantification Kit for all hESCs (N=4) and hiPSC (N=4) **(e)** Boxplot showing the median forward scatter of hESCs (N=4) and hiPSCs (N=6). **(f)** Boxplot showing the median side scatter of hESCs (N=4) and hiPSCs (N=6). **(g)** Boxplot showing the median percentage of cells across cell cycle stages for hESCs (N=4) and hiPSCs (N=6). For all boxplots, the bottom and top hinges represent the 1st and 3rd quartiles. The top whisker extends from the hinge to the largest value no further than 1.5 3 IQR from the hinge; the bottom whisker extends from the hinge to the smallest value at most 1.5 3 IQR of the hinge.

This is not the case for the results produced with the “proteomic ruler”^20^. The copy number-based analysis enables an approximation to absolute protein abundance and can reveal changes in cell mass, as we previously reported^22,23^. Using the proteomic ruler method highlighted systematic differences between hESCs and hiPSCs (Fig. 2b; Supplemental Table 3), with 56% (4,426/7,878) of all proteins detected significantly increased in hiPSCs (FC>1.5-fold; q-value < 0.001) and with particular enrichment in translation-related processes (Fig. S2). In contrast, only 40 proteins (0.5%) showed significantly lower expression levels in hiPSCs. With thousands of proteins displaying higher abundance, we hypothesised that hiPSCs have higher total protein content, compared to hESCs. Using the protein copy numbers to estimate the total protein content showed that hiPSCs had >50% higher protein content compared to hESCs (Fig. 2c). To validate this observation, an independent assay (EZQ^TM^ assay; see methods), was used to measure the total protein yield from similar numbers of freshly grown hiPSCs and hESCs. From these experiments, the calculated protein amount per million cells was >70% higher (Fig. 2d; p-value=0.0018) in hiPSCs, relative to hESCs. We conclude that hiPSCs have a higher total protein content. Changes in protein content could potentially be linked to differences in the cell cycle profile. Hence, to test this, we used fluorescence-activated cell sorting (FACS) to study the cell cycle distribution of hESCs and hiPSCs. The FACS data showed that hiPSCs have significantly higher forward scatter (Fig. 2e), correlated to increased cell size, as well as significantly higher side scatter (Fig. 2f), correlating to increased cell granularity. However, the FACS analysis revealed no significant differences between hiPSCs and hESCs in the percentage of cells at each of the cell cycle stages (Fig. 2e). We conclude that hiPSCs have significantly higher total protein content, with increased size and granularity, but that these differences with hESCs are independent of changes in cell cycle distribution.

### hiPSCs have elevated nutrient transporters, metabolic proteins, and protein and lipid synthesis machinery

To maintain a higher protein content than hEScs with a comparable cell cycle profile, hiPSCs would require higher protein synthesis capacity, which in turn requires increased access to nutrients and energy. Energy metabolism in primed pluripotent stem cells is largely dependent on glycolysis^24^, which is sensitive to glucose uptake and lactate shuttling. Therefore, we compared the expression of the respective glucose and lactate transporters between hiPSCs and hESCs. The data showed both main glucose transporters, GLUT1 (SLC2A1) and GLUT3 (SLC2A3), had higher abundance in hiPSCs, as did the lactate transporters SLC16A1 and SLC16A3 (Fig. 3a). Other rate limiting enzymes, including Hexokinase 1 (HK1) and 2 (HK2) were also significantly increased within hiPSCs (Fig. 3b), suggesting increased glycolytic potential.

**Figure 3-.**
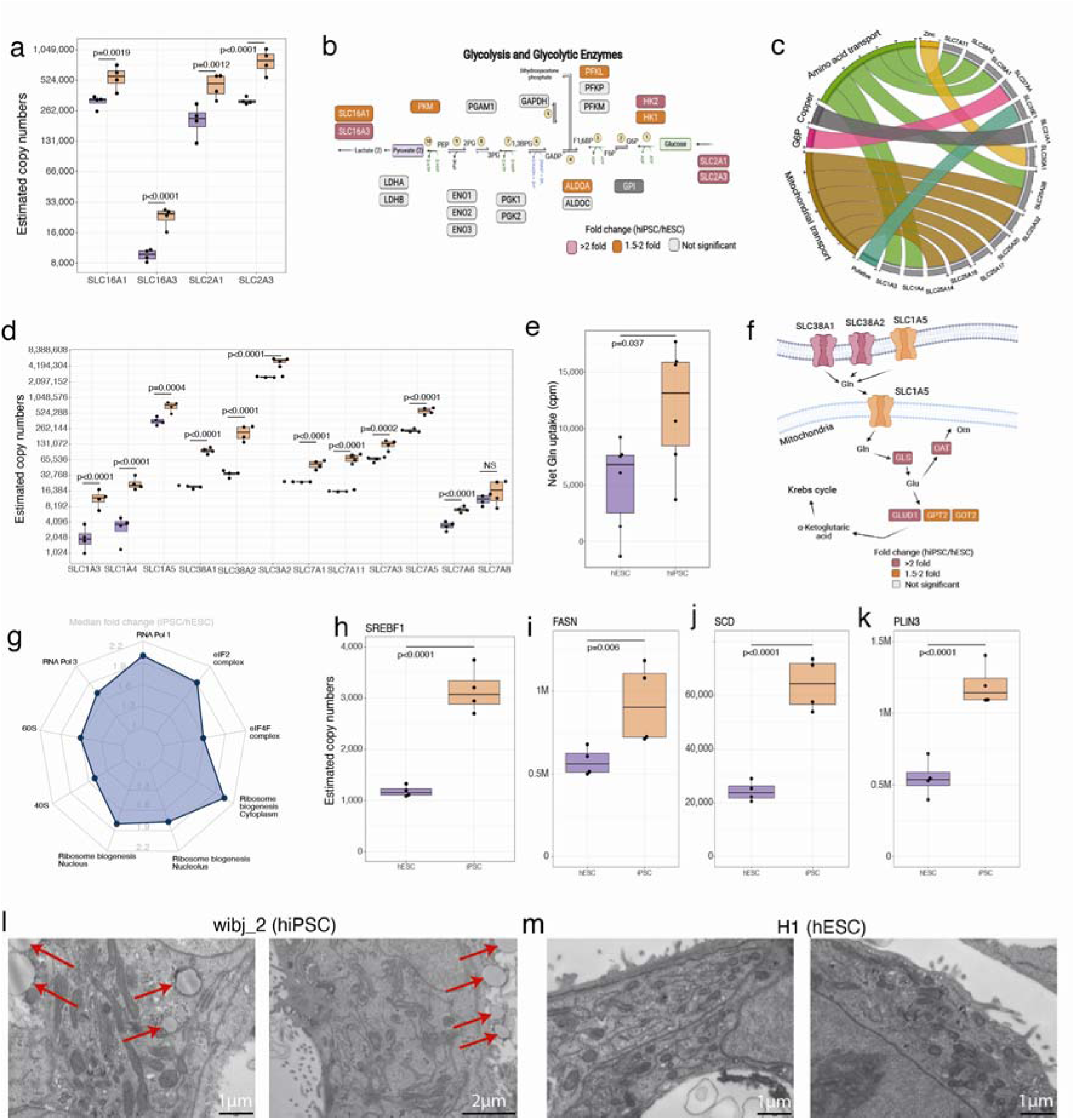
Fuelling growth: **(a)** Boxplots showing the estimated copy numbers for the lactate (SLC16A1 and SLC16A3) and glucose transporters (SLC2A1 and SLC2A3) across hESCs (N=4) and hiPSC (N=4). **(b)** Schematic showing the glycolytic proteins and their fold change in hESC vs hiPSCs. **(c)** Chord diagram showing the 15 most upregulated solute carrier proteins along with their classification based on transport activities/localisation. **(d)** Boxplots showing the estimated copy numbers of the main amino acid transporters in hESCs (N=4) and hiPSC (N=4). **(e)** Boxplot showing the net glutamine uptake (see methods) in hESCs (N=4) and hiPSC (N=4). **(f)** Schematic showing the glutaminolysis proteins and their fold change in hESCs (N=4) and hiPSC (N=4). **(g)** Radar plot showing the median fold change (hiPSC/ESC) for protein categories which are related to the pre-ribosomes. Boxplots showing the estimated copy numbers for **(h)** SREBF1, **(i)** FASN, **(j)** SCD, **(k)** PLIN3. in hESCs (N=4) and hiPSC (N=4). **(l)** Transmission electron microscopy images for wibj_2 (hiPSC). Lipid droplets are marked with red arrows. **(m)** Transmission electron microscopy images for H1 (hESC). For all boxplots, the bottom and top hinges represent the 1st and 3rd quartiles. The top whisker extends from the hinge to the largest value no further than 1.5 3 IQR from the hinge; the bottom whisker extends from the hinge to the smallest value at most 1.5 3 IQR of the hinge.

Nutrient uptake is mostly handled by the SLC (solute carrier), group of membrane transporters. Analysis of the 15 SLC transporters that are most upregulated in hiPSCs, compared to hESCs, showed that they mostly belonged to two categories, i.e., amino acid and mitochondrial transporters (Fig. 3c). Amino acids are vital to sustain high rates of protein synthesis^25^ and the data showed that 11/12 amino acid transporters were significantly increased in hiPSCs, compared to hESCs, including the hyper abundant protein SLC3A2, which is present at >4 million copies per cell (Fig. 3d). The highest fold increases, (>4-fold), were seen for SLC38A1 and SLC38A2, both of which are major glutamine transporters^26,27^. We next examined whether the increased abundance of the glutamine transporters had phenotypic impact, i.e., whether it correlated with increased glutamine uptake within hiPSCs. To test this hypothesis, we measured the uptake of radio-labelled glutamine in both hiPSCs and hESCs (see methods). The data showed that hiPSCs had a median of >90% higher uptake of glutamine, compared to hESCs (Fig. 3e). Glutamine has been reported to be the most consumed amino acid in hESCs^28^ and its catabolism to be one of the vital metabolic pathways that can provide ATP and more importantly biosynthetic precursors, required to sustain growth^29^. Hence, we also explored the abundance of enzymes involved in glutaminolysis and found that vital proteins, including GLS, GLUD1, GPT2 and GOT2, were also significantly higher in hiPSCs (Fig. 3f).

Having established that hiPSCs have increased expression of nutrient transporters and higher expression of enzymes in key metabolic pathways, compared with hESCs, we next looked at the machinery required for protein synthesis. The levels of many of the proteins involved in ribosome subunit biogenesis, including ribosomal proteins, were higher in hiPSCs (Fig. 3g). The increased expression of translation machinery components, nutrient transporters and many metabolic enzymes, is consistent with the increased total protein content seen within hiPSCs.

The data also highlighted increased fatty acid (FA) and lipid droplet (LD) synthesis potential in hiPSCs, with increased abundance of the protein SREBP1 (SREBF1; Fig. 3h), a master regulator of lipid synthesis^30^, as well as FASN (Fig. 3i) and SCD (Fig. 3j). Similarly, a crucial regulator for LD assembly, PLIN3^31^, (Fig. 3k), displayed >2-fold increased abundance in hiPSCs. To examine the potential phenotypic impact of this increased abundance of proteins involved in LD synthesis, we performed transmission electron microscopy (TEM) analyses to compare hiPS and hES cells. This showed that LDs were clearly visible in hiPSCs (Fig. 3l), but not visible in hESCs (Fig. 3m). We conclude that the hiPSCs have elevated levels of LDs, resulting from the increased expression of proteins involved in lipid synthesis and LD assembly.

### hiPSCs show altered mitochondrial metabolism compared to hESCs

Our data also highlighted important changes in mitochondrial proteins, including increases in the levels of metabolic proteins that are encoded within the mitochondrial genome^32^, (Fig. 4a). The latter proteins, which are translated by specialised mitochondrial ribosomes (mitoribosomes), are embedded in the mitochondrial membrane. The protein components of mitoribosomes also showed increased expression in hiPSCs (Fig. 4b), along with virtually all proteins involved in the translation initiation, elongation and termination of mitochondrial genome-encoded proteins (Fig. 4c).

**Figure 4-.**
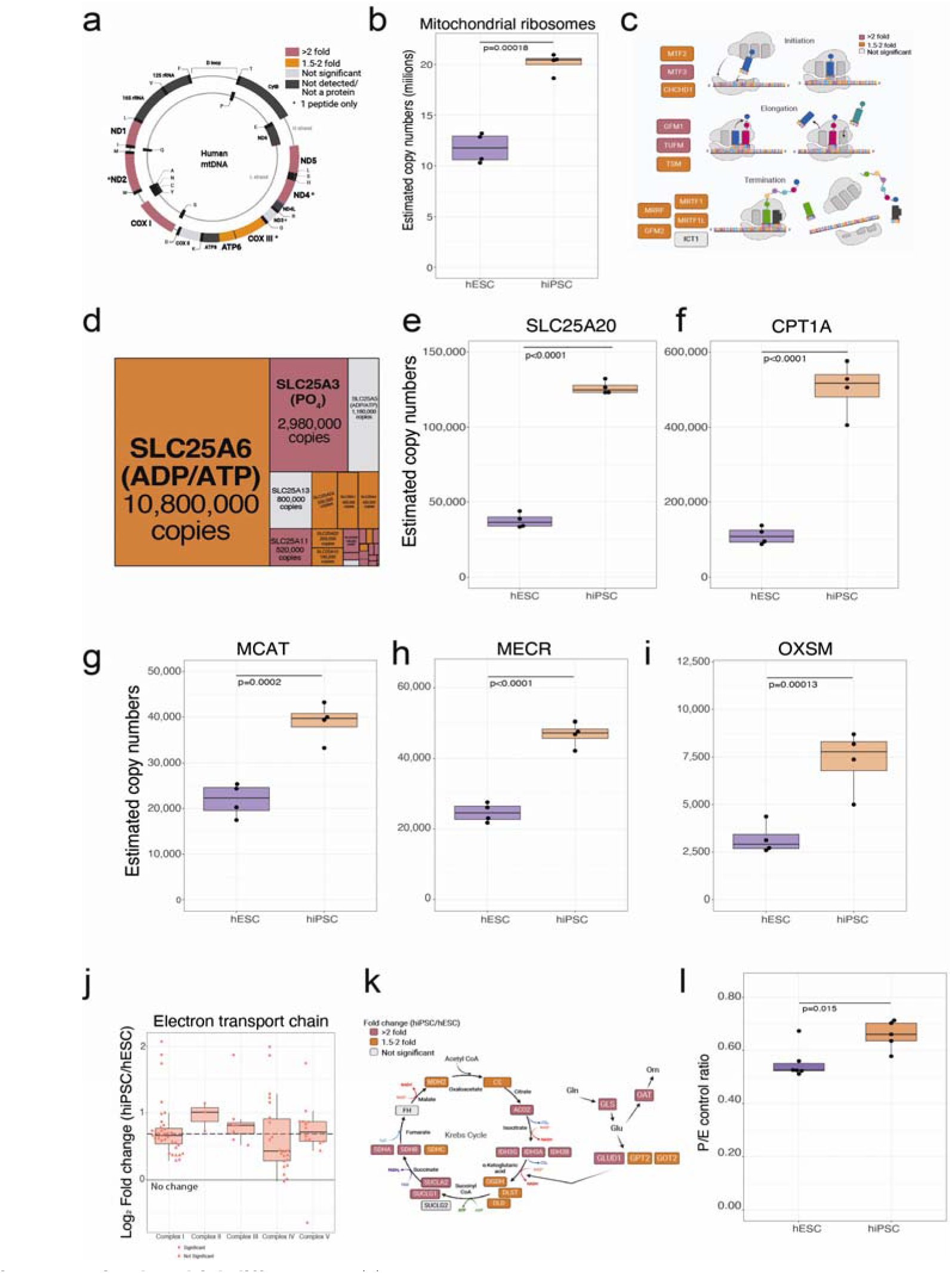
Mitochondrial differences: **(a)** Schematic showing the mitochondrial genome encoded proteins and their fold change in hESCs and hiPSCs. **(b)** Boxplot showing the estimated copy numbers of all mitochondrial ribosomal proteins hESCs (N=4) and hiPSC (N=4)**(c)** Schematic showing proteins involved in mitochondrial translation and their fold change (hiPSCs/hESCs). **(d)** Treeplot showing all mitochondrial transporters, size is proportional to the median estimated copy numbers in hiPSCs (N=4). Boxplot showing the estimated copy numbers for **(e)** SLC25A20. **(f)** CPT1A. **(g)** MCAT, **(h)** MECR, **(i)** OSXM, in hESCs (N=4) and hiPSC (N=4)**(j)** Boxplot showing the log2 fold change (hiPSC/hESCs) for all subunits of the different complexes of the electron transport chain. The median fold change across all detected proteins is shown as a dotted line. **(k)** Schematic showing the fold change of critic acid cycle and glutaminolysis proteins in hiPSCs (N=4) vs hESCs (N=4). **(l)** Boxplot showing the P (oxphos capacity) /E (electron transfer capacity) control ratio. For all boxplots, the bottom and top hinges represent the 1st and 3rd quartiles. The top whisker extends from the hinge to the largest value no further than 1.5 3 IQR from the hinge; the bottom whisker extends from the hinge to the smallest value at most 1.5 3 IQR of the hinge.

The analysis of transporter proteins revealed a cluster of 22/27 mitochondrial transporters that were significantly increased in hiPSCs, including the hyper abundant, (>10 million copies per cell), ATP/ADP transporter (Fig. 4d). A subset of 14 transporters displayed >2-fold increased abundance, including the acylcarnitine transporter SLC25A20 (Fig. 4e), which is part of the carnitine shuttle in the beta oxidation pathway. Another component of the carnitine shuttle, CPT1A, displayed over 4-fold higher abundance in hiPSCs (Fig. 4f), suggesting an important role. The data showed it was not just fatty acid oxidation, but also synthesis, that was affected, with proteins acting in the mitochondrial fatty acid synthesis (mFAS) pathway also increased in abundance. MCAT (Fig. 4g), MECR (Fig. 4h) and OXSM (Fig. 4i), all displayed ~2-fold higher abundance in hiPSCs compared to hESCs. These results have a metabolic relevance as mFAS has been reported to control the activity of the electron transport chain (ETC) ^33^, which was also increased in hiPSCs, with subunits of all 5 ETC complexes increased in abundance in hiPSCs and with complex II and complex III showing the most prominent effects (Fig. 4j). Complex II is also part of the tricarboxylic acid (TCA) cycle, which displayed increased abundance of the majority of proteins involved in the pathway (Fig. 4k).

As the proteomic data showed clear differences between hiPSCs and hESCs in the levels of mitometabolism proteins, we performed experiments to explore whether this was reflected in phenotypic differences between hiPSCs and hESCs. This was tested using high-resolution respirometry (see Methods). The data showed that hiPSCs had a higher P/E control ratio to hESCs, which denotes an increased capacity of the phosphorylation system to produce ATP (Fig. 4l). We conclude that hiPSCs have elevated levels of mitometabolism proteins relative to hESCs, resulting in higher respiratory activity.

### hiPSCs upregulate secreted proteins affecting their microenvironment

Among the most upregulated proteins in hiPSCs were a subset of secreted proteins. Secreted proteins are of great importance because changes in their absolute abundance can affect the extracellular environment. These secreted proteins mostly represented 4 categories: structural extracellular matrix (ECM) proteins, growth factors, protease inhibitors and proteases (Fig. 5a). The ECM both provides physical support for cells, as well as actively participating in cell signalling by providing domains for growth factors^34^. The ECM can also be reshaped in tumours, thereby promoting cancer cell growth and migration^35^. The current data show that both laminins and collagens were all increased in abundance in hiPSCs (Fig. 5b). Collagens are reported to alter the stiffness of the ECM and their synthesis is iron intensive. Interestingly, the data also show that proteins involved in importing and storing iron were increased in abundance in hiPSCs (Fig. 5c-f).

**Figure 5-.**
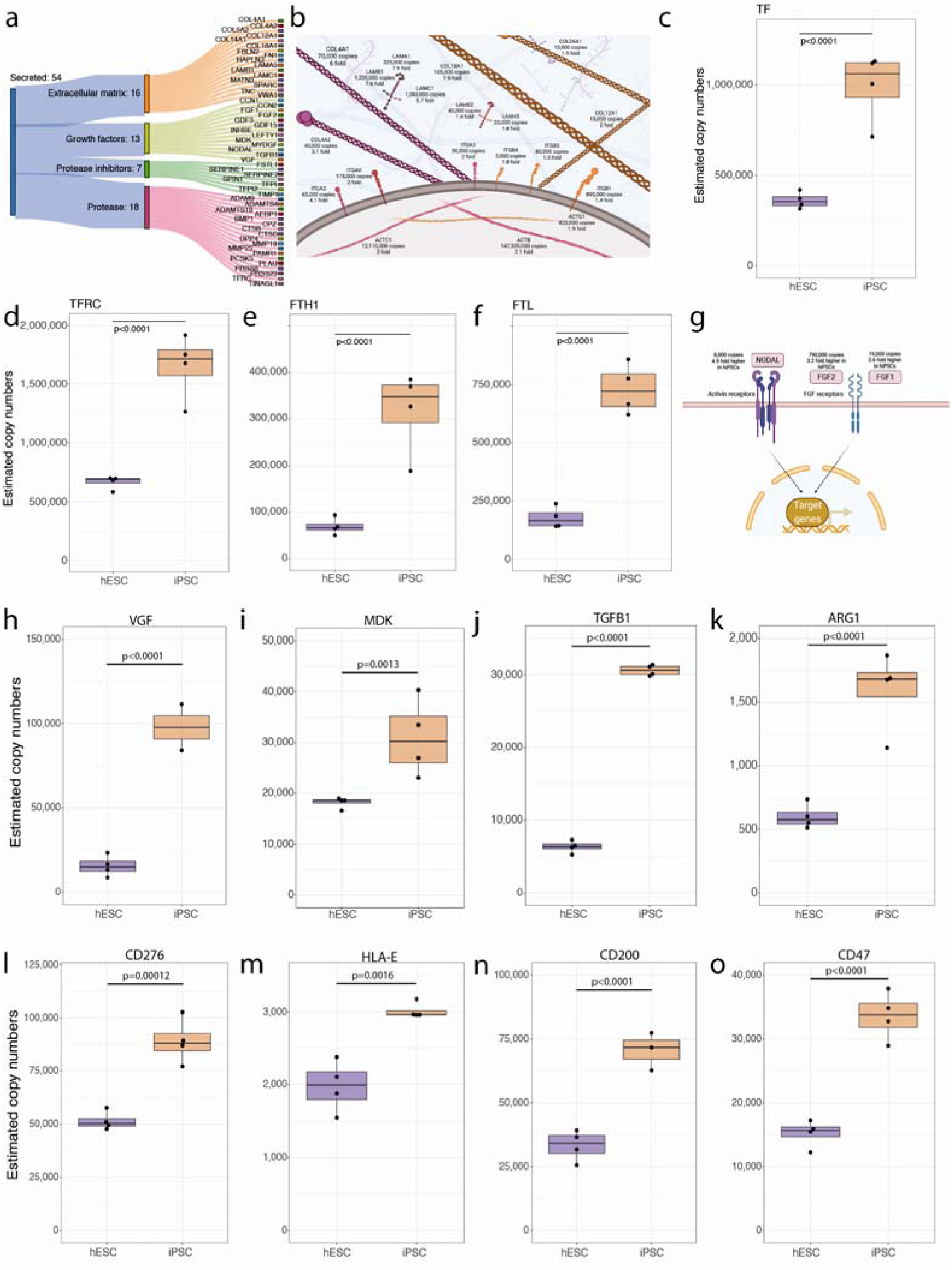
Secreted proteins: **(a)** Sankey diagram showing the secreted proteins that belong to the extracellular matrix (ECM), Growth factor, Protease Inhibitor or Protease categories and are significantly increased in abundance in hiPSCs **(b)** Schematic showing ECM proteins that are significantly increased in abundance in hiPSCs. Boxplot showing the estimated copy numbers for (**c)** TF, **(d)** TFRC, **(e)** FTH1, and **(f)** FTL in hESCs (N=4) and hiPSC (N=4). **(g)**Schematic showing the changes in abundance in primed pluripotency growth factors. **(h)** Boxplot showing the estimated protein copy numbers for VGF. **(i)** Boxplot showing the estimated protein copy numbers for MDK. Boxplot showing the estimated protein copy numbers for **(j)** TGFB1, **(k)** ARG1, **(l)** CD276, **(m)** HLA-E, **(n)** CD200 and **(o)** CD47. All boxplots show the data for hESCs and hiPSCs. For all boxplots, the bottom and top hinges represent the 1st and 3rd quartiles. The top whisker extends from the hinge to the largest value no further than 1.5 3 IQR from the hinge; the bottom whisker extends from the hinge to the smallest value at most 1.5 3 IQR of the hinge.

The data also showed that 13 growth factors were increased in abundance in hiPSCs, compared to hESCs. A subset of these, i.e., FGF1, FGF2 and NODAL, are reported to have direct relevance to the maintenance of pluripotency and can modulate important processes in PSCs ^36–38^ (Fig. 5g). Other growth factors that are upregulated in hPSCs are linked to disease mechanisms and cancer. This includes VGF (Fig. 5h), which is linked to promoting growth and survival in glioblastoma^39^ and MDK (Fig. 5i), which is highly expressed in malignant tumors^40^ and has been shown to play a role in chemoresistance^41^.

### hiPSCs display increased abundance of immunosuppressive proteins

NODAL wasn’t the only growth factor in the TGFB family that was increased in hiPSCs, with TGFB1 displaying a ~5-fold increase in abundance in hiPSCs compared to hESCs (Fig. 5j). Besides its role as a growth factor, TGFB1 has been shown to have important roles in the regulation of the immune response, promoting the generation of regulatory T cells, while inhibiting the generation and function of effector T cells. As immunogenicity of PSCs is a topic of relevance to clinical adaptations, we looked for differences in modulators of the immune response.

Arginine availability is vital to effector T cells and other leukocytes, where depletion mediated by Arginase has been shown to be linked to T cell inhibition^42^. Our data show that hiPSCs have ~2.5-fold higher abundance of ARG1 (Fig. 5k). Furthermore, hiPSCs also display increased expression of the immune checkpoint protein, CD276 (Fig. 5l), which has been reported to be a potent inhibitor of survival and function of T cells^43,44^.

hiPSCs also displayed increased abundance of inhibitory ligands that supress the immune function of other leukocytes. The data show hiPSCs have increased abundance of the non-classical HLA-E (Fig. 5m), which has been shown to interact with the NK cell receptor, NKG2A, to mediate immune evasion in ageing cells^45^. They also displayed increased abundance of CD200 (Fig. 5n), a ligand for CD200R, which can inhibit the immune response from macrophages, basophils, NK cells and T cells, as well as CD47 (Fig. 5o), a ligand of SIRPA that helps cells to escape macrophage phagocytosis. These data indicate that hiPSCs have increased abundance of known immunosuppressive proteins, compared to hESCs.

### hiPSCs display reduced abundance of H1 histones

A striking feature of this proteomic study is how few proteins (<1%; 40/7,878), showed significantly decreased abundance in hiPSCs, compared to hESCs. A high proportion of these decreased abundance proteins affect nuclear processes. Thus, an overrepresentation analysis showed that proteins whose abundance was decreased in hiPSCs were enriched in GO terms related to DNA recombination, nucleosome positioning and chromatin silencing (Fig. 6a). Notably, this included four H1 histone variants, which are reported to influence nucleosomal repeat length^46^ and stabilise chromatin structures^47^. Our data show that the most abundant H1 variant in hESCs, HIST1H1E, is decreased in abundance in hiPSCs by ~3.5-fold (Fig. 6b), while HIST1H1C (Fig. 6c), HIST1H1D (Fig. 6d) and H1FX (Fig. 6e) are all decreased by >1.7-fold.

**Figure 6 –.**
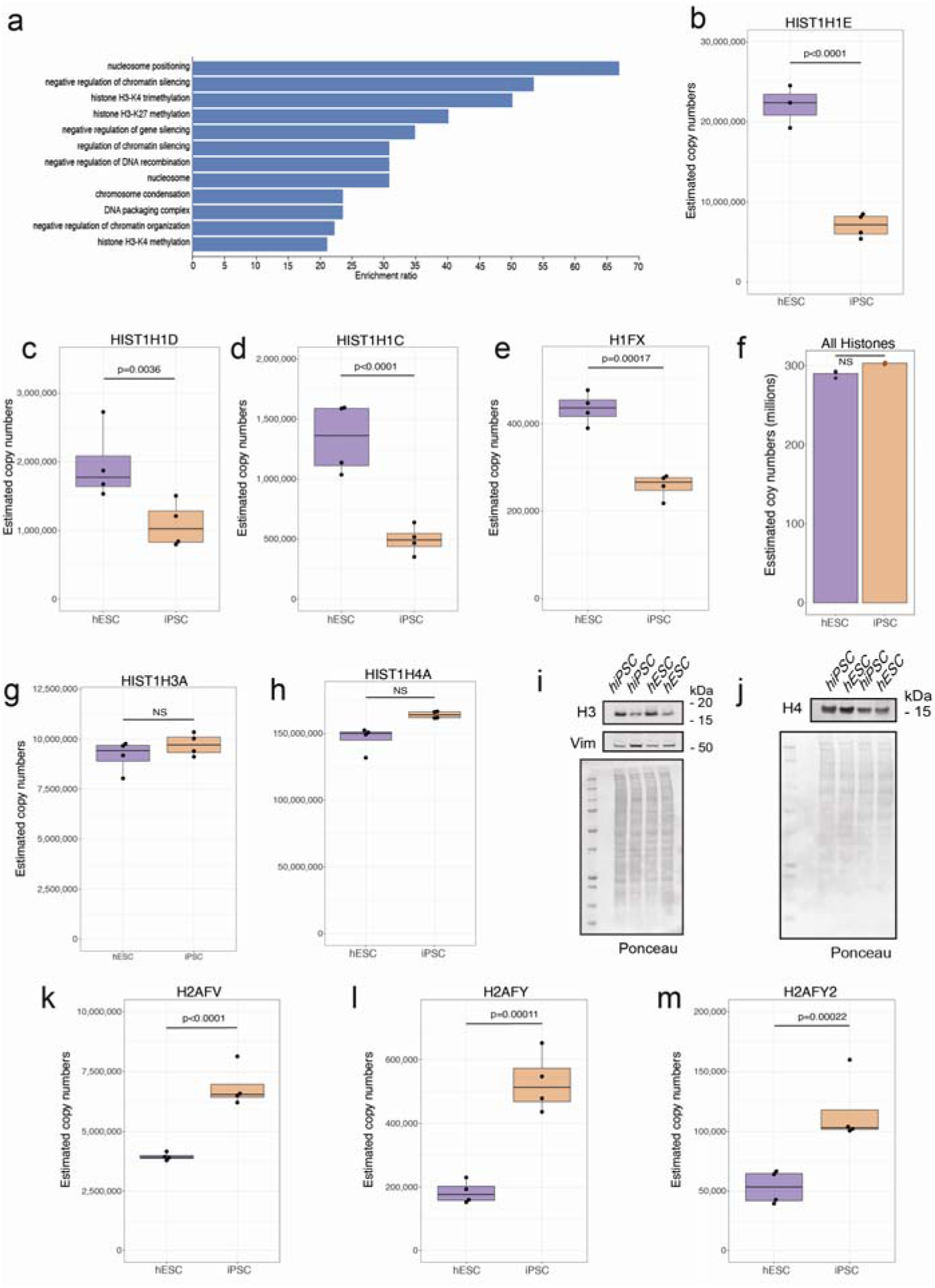
Changes within histones: **(a)** Barplot showing the GO term enrichment results for proteins significantly decreased in abundance (see methods) in hiPSCs. Boxplots showing estimated copy numbers for **(b)** HIST1H1E, **(c)** HIST1H1D, **(d)** HIST1H1C and **(e)** H1FX in hESCs (N=4) and hiPSC (N=4). **(f)** Barplot showing the median estimated copy numbers for all histones in hESCs and hiPSCs. Boxplots showing estimated copy numbers for **(g)** HIST1H3A in and **(h)** HIST1H4A in hESCs and hiPSCs. Western blot showing the abundance of **(i)** H3 and **(j)** H4 histones in hESCs (N=4) and hiPSC (N=4). Boxplots showing estimated copy numbers for **(k)** H2AFV, **(l)** H2AFY and **(m)** H2AFY2 in hESCs (N=4) and hiPSC (N=4). For all boxplots, the bottom and top hinges represent the 1st and 3rd quartiles. The top whisker extends from the hinge to the largest value no further than 1.5 3 IQR from the hinge; the bottom whisker extends from the hinge to the smallest value at most 1.5 3 IQR of the hinge.

As histone variants have very similar protein sequences, where peptides detected by MS can potentially match to multiple H1 histone variants, a peptide level analysis was necessary to deconvolute the signal (Fig. S3). The Andromeda search engine^48^ assigns peptide intensities to a protein following a razor peptide approach, where the intensity of a peptide is assigned to only one protein, regardless of whether it is unique, or shared by two or more proteins. This makes the analysis of specific variants challenging at the protein level. Hence, we focussed on a peptide-specific approach with the MS data and found that the intensity of the peptides that were shared between these 3 H1 histone variants displayed a consistent reduction in abundance in hiPSCs (Fig. S3).

The systematic reduction in abundance in hiPSCs seen with H1 histone variants was not seen for members of the other histone families. Evaluating either the concentration (Fig. S1), or copy numbers (Fig. 6f), across all histones, showed no significant differences in expression between hiPSCs and hESCs. Furthermore, for core histones, including H3 and H4, there were no significant abundance differences seen within either the proteomics data (Fig. 6g&h), or in additional western blot analyses that were performed to validate these conclusions (Fig. 6i&j). However, we did detect differences between hiPSCs and hESCs in the expression of histone H2 variants, with H2AFV (Fig. 6k), H2AFY (Fig. 6l) and H2AFY2 (Fig. 6m), all increased in abundance in hiPSCs. As histone H2 variants also have high sequence similarity and shared peptides, we also performed a peptide level analysis, which validated that both shared and unique peptides displayed the same pattern, i.e., showing increased abundance in hiPSCs (Fig. S4). Thus, we conclude that there are opposing effects for histone H1 and histone H2 variants, with the former decreased and the latter increased in abundance in hiPSCs.

## Discussion

Induced pluripotent stem cells can provide valuable models for clinical research and future therapies, which makes it vital to understand both their similarities and any specific differences, with embryo-derived human stem cells. This study provides a detailed comparison of the proteomes of multiple hiPSC and hESC lines derived from different donors. The major conclusion is that while hiPSC and hESC lines express a near identical set of proteins, with similar abundance ranks, they also display important quantitative differences. In particular, our data indicate that hiPSCs reprogrammed from skin fibroblasts display considerable differences in their cytoplasmic and mitochondrial proteomes, compared to hESCs, while the nuclear proteome was very similar between the two cell types. Furthermore, additional microscopy analyses and functional assays showed that the systematic differences in the proteomes of the respective hiPSCs and hESCs had a measurable impact on cell phenotypes, most notably affecting mitochondria, metabolic activity, and nutrient transport. It would be of interest in future to analyse whether hiPSCs reprogrammed from different cell types, such as peripheral leukocytes, also share these differences in protein expression with hESCs, or if these specific changes are characteristic of fibroblast-derived iPS cells.

Using estimated protein copy numbers, our data show that only <1% of proteins were significantly decreased in abundance in hiPSCs, compared to hESCs, including multiple H1 histones In contrast, ~ 56% of all proteins quantified were significantly increased in hiPSCs (fold change>1.5 and q-value <0.001), with most of these increases affecting cytoplasmic and mitochondrial proteins and activities. The MS data show that total protein levels are higher overall in hiPSCs, as compared with hESCs, a result that was independently validated using an EZQ protein assay. This difference in total protein content was shown by FACS analysis not to result from systematic differences between hiPSCs and hESCs in cell cycle progression. Instead, the increased protein levels in hiPSCs correlated with increased levels of the protein translational machinery, along with increased metabolic and mitochondrial activity and higher levels of nutrient transport.

These results highlight an important technical point relating to data normalisation and its effect on the interpretation of such data. By using a standard median normalisation (concentration-based approach), instead of the proteomic ruler^20^, the difference in total protein content between the cell types, involving the increased abundance of thousands of proteins, is not apparent. Hence, two cell types with, for example, a 4-fold higher total protein content, could nonetheless appear to show virtually no significant differences, as long as the protein ranks and relative concentrations remained similar. This would result in an erroneous conclusion that there is little to no change in protein expression, while the orthogonal data suggest otherwise.

Having established that hiPSCs displayed higher total protein content than corresponding hESC lines, we sought to understand how this could be maintained. To maximise protein synthesis, nutrient availability and energy production are key factors^25^. The proteomic data show that vital nutrient transporters, known to be important for growth and protein production,^49^ were significantly increased in hiPSCs, compared to hESCs. In particular, the 3 glutamine transporters, SLC1A5, SLC38A1 and SLC38A2, were all significantly increased in abundance, with additional functional assays showing that this correlated with higher levels of glutamine uptake measured in hiPSCs. Glutamine has been previously shown to fuel growth and proliferation in rapidly dividing cells, including cancer cells^26^ and could be sustaining higher rates in hiPSCs.

Nutrients provide the fuel, but it is the metabolic proteins that are the engines that convert them to energy. Here, our data showed that proteins involved in both glycolysis and glutaminolysis were significantly increased in abundance in hiPSCs. When cells preferentially use the glycolytic pathway, e.g., stem cells and cancer cells, there is increased demand for biosynthetic precursors and NADPH^50^. These precursors can be supplied via glutaminolysis^51–53^ linked to the TCA, both important mitochondrial processes and both with significantly increased protein levels in hiPSCs, along with other proteins involved in the electron transport chain. One caveat is that an uneven distribution of biological sex between donors of the respective hESC and hiPSCs lines, could mean there is a potential for sex-specific differences to explain at least some of these effects, however in previous a large-scale proteomic study of male and female hiPSCs no changes in proteins involved in oxidative phosphorylation were detected^54^. Differences in the mitochondria between hiPSCs and hESCs have been previously reported, but whether they originate from the reprogramming process, or are induced by the increased nutrient uptake, remains unknown and is a point of interest for future studies.

Secreted proteins, such as growth factors and ECM proteins, are a category of great interest, because their absolute abundance can affect the surrounding cellular microenvironment. hiPSCs were found here to show increased expression levels of growth factors that are linked to cancer and immunosuppression. For example FGF2, an important growth factor for primed pluripotent stem cells, has been shown that to promote ERK activation^55^, stimulating protein synthesis^56–58^. Thus, the increased abundance of FGF2 could generate a feedforward loop, further driving/sustaining growth in hiPSCs, however that growth potential is also linked to breast^59^ and gastric cancers^60^, as well as gliomas^61^. Another important growth factor that is increased in abundance in hiPSCs is TGFB1, a known potent inhibitor of T cell responses^62,63^. We note that the immunogenicity of pluripotent stem cells has important consequences for cell therapy applications. Our data suggest that hiPSCs might have a higher immune evasion potential, via multiple mechanisms. They display increased abundance of secreted T cell inhibitors, including TGFB1 and ARG1^64^, along with inhibitory ligands, such as CD276, CD200 and CD47. An increased inhibitory capacity, combined with the tumorigenic potential of hiPSCs^65^, raises some concerns about the suitability of using reprogrammed hiPSCs for certain types of therapeutic applications. Based upon our current data, we recommend that potential phenotypic consequences resulting from the observed differences in iPS and ESC proteomes should be studied further to determine whether the immunosuppressive properties of these cells vary significantly.

In summary, our data show that hiPSCs and hESCs, despite their clear similarities, are not identical at both the protein and phenotypic levels. We show that hiPSCs reprogrammed from skin fibroblasts differ from hESCs, predominantly in their cytoplasmic and mitochondrial proteome, leading to measurable functional differences affecting their metabolic activity and growth potential. These data can help to inform future strategies to mitigate for these differences as hiPSCs continue to be used in important clinical applications and as disease models.

## Materials and Methods

### HipSci hiPSC line generation

As part of the HipSci project^16^ hiPSC lines were generated in the sanger centre. hiPSCs were generated from fibroblasts obtained by skin punch biopsies. The fibroblasts were reprogrammed as described previously^16^, in brief Sendai vectors expressing hOCT3/4, hSOX2 and hc-MYC were used.

### HipSci hiPSC line quality control

The hiPSC lines were passaged a mean of 16 times before being subjected to the first tier of molecular data for quality control, which included genotyping (‘gtarray’), gene expression data (‘gexarray’), and an assessment of the pluripotency and differentiation potential of each line (‘Cellomics’). Pluripotency of the lines was additionally verified *in silico*, using the PluriTest assay^66^. Subsequently one or two lines per donor were subjected to a set of molecular data QC assays. The criteria for line selection were: (i) level of pluripotency, as determined by the PluriTest assay (ii) number of copy number abnormalities and (iii) ability to differentiate into each of the three germ layers. These included proteomics, DNA methylation (‘mtarray’), RNA-sequencing and high-content cellular imaging.

### hiPSC and hESC Cell Culture

Human iPS cells generated by the HipSci consortium^16^ (aizi_1,bubh_3, kucg_2, oaqd_3, ueah_1 and wibj_2) and human hESCs (SA121 and SA181, H1, H9) were both grown in identical conditions, maintained in TESR medium^67^ supplemented with FGF2 (Peprotech, 30 ng/ml) and noggin (Peprotech, 10 ng/ml) on growth factor reduced geltrex basement membrane extract (Life Technologies, 10 μg/cm2) coated dishes at 37°C in a humidified atmosphere of 5% CO_2_ in air.

Cells were routinely passaged twice a week as single cells using TrypLE select (Life Technologies) and replated in TESR medium that was further supplemented with the Rho kinase inhibitor Y27632 (Tocris, 10 μM) to enhance single cell survival. Twenty-four hours after replating Y27632 was removed from the culture medium. For proteomic analyses cells were plated in 100 mm geltrex coated dishes at a density of 5×10^4^ cells cm^−2^ and allowed to grow to for 3 days until confluent with daily medium changes.

### Immunoblotting

Equal volumes of hiPSC or hESCs protein lysates were boiled in LDS/RA buffer for 5 mins at 95°C and loaded into 4-15% NuPAGE Bis-Tris SDS-PAGE gels in running buffer (50 mM MES, 50 mM Tris, 0.1 % SDS, 1 mM EDTA, pH7.3), transferred onto nitrocellulose membrane (Amersham #10600041) in transfer buffer (8 mM Tris, 30 mM Glycine, 20 % Methanol) and stained with Ponceau S (Sigma-Aldrich, #P7170). Membranes were blocked in TBS-T + 5% BSA for 1 hr at RT and incubated overnight at 4°C in primary antibodies prepared in TBS-T + 5% BSA. Membranes were washed 3 x 15 mins in TBS-T, incubated with secondary antibody for 1 hr at RT, washed, and imaged using Odyssey CLx (LI-COR). Antibodies: Histone H3 (Abcam, ab1791, 1:1000); Histone H4 (Abcam, ab10158, 1:1000), Vimentin (Cell Signalling Technology (CST), #5741S, 1:1000), IRDye® 680RD Donkey anti-Rabbit IgG Secondary Antibody (LI-COR, 926-68073, 1:10,000), GAPDH (CST, #97166, 1:10000), OCT4A (CST, #2840, 1:10000), SOX2 (CST, #3579, 1:10000), NANOG (CST, #4903, 1:10000).

### Flow cytometric analysis

Cells were seeded on geltrex coated dishes in TESR medium at a density of 3×10^3^ cells/cm^2^. After 24 hours the medium was replaced with fresh TESR medium and after a further 24 hours the cells were harvested using TrypLE select, resuspended in TESR medium and counted (cell density at harvest was approximately 1×10^5^ cells/cm^2^ for each cell line). 5×10^5^ cells were then collected by centrifugation at 300 xg for 2 minutes then resuspended with 1ml of Dulbecco’s PBS (without calcium or magnesium) containing 1% fetal bovine serum. The cells were then collected by centrifugation at 300 xg for 2 minutes and resuspended in 1ml of ice cold 90% methanol / 10% dH_2_O while vortexing. Samples were then incubated for 30 minutes at room temperature before being stored at −20°C until they were analysed.

For cell cycle analysis the cells were collected by centrifugation at 300 xg for 2 minutes, washed with PBS, then resuspended in 300 micro l of staining buffer (Dulbecco’s PBS + 1% FBS + 50 micro g/ml propidium iodide, 50 micro g/ml ribonuclease A) for 20 minutes at room temperature in the dark. Cellular DNA content was determined by analysis on a FACS Canto flow cytometer (BD Biosciences). PI fluorescence was detected using 488nm excitation and fluorescence emission collected at 585/42nm. Data was analysed using Flowjo software. Doublet discrimination was performed on the basis of PI-A v PI-W measurements and cell cycle distribution determined using the Watson Pragmatic model.

### Cell line selection for mass spectrometry

Human iPS cells (bubh_3, kucg_2, oaqd_3 and wibj_2) and human hESCs (SA121 and SA181, H1 and H9) were analysed by mass spectrometry using TMT as described below.

### Protein extraction

Cell pellets were resuspended in 300 µL extraction buffer (4% SDS in 100 mM triethylammonium bicarbonate (TEAB), phosphatase inhibitors (PhosSTOP™, Roche)). Samples were boiled (15 min, 95 °C, 350 rpm) and sonicated for 30 cycles in a bath sonicator (Bioruptor® Pico bath sonicator, Diagenode, Belgium; 30s on, 30s off) followed by probe sonication for 50 s (20s on, 5s off). 2 µL Benzonase® nuclease HC (250 U/µL, Merck Millipore) was added and incubated for 30 min (37 °C, 750 rpm). Reversibly oxidized cysteines were reduced with 10 mM TCEP (45 min, 22 °C, 1,000 rpm) followed by alkylation of free thiols with 20 mM iodoacetamide (45 min, 22 °C, 1,000 rpm, in the dark). Proteins were quantified using the fluorometric EZQ^TM^ assay (Thermo Fisher Scientific).

### Protein digestion using the SP3 method

Protein extracts were cleaned and digested with the SP3 method as described previously with modifications^68,69^. Briefly, 50 µL of a 20 µg/µL SP3 bead stock (Sera-Mag SpeedBead carboxylate-modified magnetic particles; GE Healthcare Life Sciences) and 500 µL acetonitrile (ACN; final concentration of 70%) were added to 150 µL of protein extract and incubated at room temperature for 10 min (1,000 rpm). Tubes were mounted on a magnetic rack, supernatants were removed and beads were washed twice with 70% ethanol and once with ACN (1 mL each). Beads were resuspended in 80 µL 100 mM TEAB and digested for 4 h with LysC followed by tryptic digestion overnight (1:50 protease:protein ratio, 37 °C, 1,000 rpm). Peptides were cleaned by addition of 3.5 µL formic acid (final concentration of 4%) and 1.7 mL ACN (final concentration of 95%) followed by incubation for 10 min. After spinning down (1,000 g) tubes were mounted on a magnetic rack and beads were washed once with 1.5 mL ACN. Peptides were eluted from the beads with 100 µL 2% DMSO and acidified with 5.2 µL 20 % formic acid (final concentration of 1%) followed by centrifugation (15,000 g). Peptide amounts were quantified using the fluorometric CBQCA assay (Thermo Scientific).

### TMT labelling

For each sample 15 µg peptides per sample were dried *in vacuo* in a Concentrator plus (Eppendorf) and resuspended in 50 µL 200 mM EPPS pH 8.5. TMT10plex tags (Thermo Scientific) were dissolved in anhydrous ACN and added to the peptide sample in a 1:10 peptide:TMT ratio. Additional anhydrous ACN was added to a final volume of 22 µL. Samples were incubated for 2 h (22 °C, 750 rpm). Unreacted TMT was quenched by incubation with 5 µL 5% hydroxylamine for 30 min. Samples were combined, dried *in vacuo* and resuspended in 1% TFA followed by clean-up with solid-phase extraction using Waters Sep-Pak tC18 50 mg. Samples were loaded, washed five times with 1 mL 0.1% TFA in water and peptides were eluted with 70% ACN/0.1% TFA (1 mL) and dried *in vacuo* in a Concentrator plus (Eppendorf).

### High pH reversed phase peptide fractionation

TMT labelled peptide samples were fractionated using off-line high pH reverse phase chromatography. Dried samples were resuspended in 5% formic acid and loaded onto a 4.6 x 250 mm XBridge BEH130 C18 column (3.5 µm, 130 Å; Waters). Samples were separated on a Dionex Ultimate 3000 HPLC system with a flow rate of 1 mL/min. Solvents used were water (A), ACN (B) and 100 mM ammonium formate pH 9 (C). While solvent C was kept constant at 10%, solvent B started at 5% for 3 min, increased to 21.5% in 2 min, 48.8% in 11 min and 90% in 1 min, was kept at 90% for further 5 min followed by returning to starting conditions and re-equilibration for 8 min. Peptides were separated into 48 fractions, which were concatenated into 24 fractions and subsequently dried *in vacuo*. Peptides were redissolved in 5% formic acid and analysed by LC-MS.

### LC-MS analysis

TMT labelled samples were analysed on an Orbitrap Fusion Tribrid mass spectrometer coupled to a Dionex RSLCnano HPLC (Thermo Scientific). Samples were loaded onto a 100 µm × 2 cm Acclaim PepMap-C18 trap column (5 µm, 100 Å) with 0.1% trifluoroacetic acid for 7 min and a constant flow of 4 µL/min. Peptides were separated on a 75 µm × 50 cm EASY-Spray C18 column (2 µm, 100 Å; Thermo Scientific) at 50 °C using a linear gradient from 10% to 40% B in 153 min with a flow rate of 200 nL/min. Solvents used were 0.1% formic acid (A) and 80% ACN/0.1% formic acid (B). The spray was initiated by applying 2.5 kV to the EASY-Spray emitter. The ion transfer capillary temperature was set to 275 °C and the radio frequency of the S-lens to 50%. Data were acquired under the control of Xcalibur software in a data-dependent mode. The number of dependent scans was 12. The full scan was acquired in the orbitrap covering the mass range of *m/z* 350 to 1,400 with a mass resolution of 120,000, an AGC target of 4×10^5^ ions and a maximum injection time of 50 ms. Precursor ions with charges between 2 and 7 and a minimum intensity of 5×10^3^ were selected with an isolation window of *m/z* 1.2 for fragmentation using collision-induced dissociation in the ion trap with 35% collision energy. The ion trap scan rate was set to “rapid”. The AGC target was set to 1×10^4^ ions with a maximum injection time of 50 ms and a dynamic exclusion of 60 s. During the MS3 analysis, for more accurate TMT quantification, 5 fragment ions were co-isolated using synchronous precursor selection in a window of *m/z* 2 and further fragmented with a HCD collision energy of 65%. The fragments were then analysed in the orbitrap with a resolution of 50,000. The AGC target was set to 5×10^4^ ions and the maximum injection time was 105 ms.

### TMT channel allocation

| TMT tag | Cell line |
| --- | --- |
| 126C | bubh_3 (hiPSC - female) |
| 127N | H01 (hESC - male) |
| 127C | kucg_2 (hiPSC - male) |
| 128N | H09 (hESC - female) |
| 128C | oaqd_3 (hiPSC - male) |
| 129N | SA121 (hESC - male) |
| 129C | wibj_2 (hiPSC - female) |
| 130N | SA181 (hESC - male) |
| 130C | Empty |
| 131 | Empty |

### High-resolution respirometry in wibj_2 and H1 stem cells

Mitochondrial respiration was studied in digitonin-permeabilised wibj_2 and H1 stem cells (10 μg / 1×10^6^ cells), to keep mitochondria in their architectural environment. The analysis was performed in an oxygraphic chamber with thermostat set to 37°C with continuous stirring (Oxygraph-2 k, Oroboros instruments, Innsbruck, Austria). Cells were collected with trypsin, pelleted, and then placed in MiR05 respiration medium (110 mM sucrose, 60 mM lactobionic acid, 0.5 mM EGTA, 3 mM MgCl_2_, 20 mM taurine, 10 mM KH_2_PO_4_, 20 mM HEPES adjusted to pH 7.1 with KOH at 30°C, and 1 g/l BSA, essentially fatty acid free). Substrate-Uncoupler-Inhibitor titration protocol number 2 (SUIT-002)^70^ was used to determine respiratory rates. Briefly, after residual oxygen consumption in absence of endogenous fuel substrates (ROX, in presence of 2.5 mM ADP), was measured, fatty acid oxidation pathway state (F) was evaluated by adding malate (0.1 mM) and octanoyl carnitine (0.2 mM) (OctM_P_). Membrane integrity was tested by adding cytochrome c (10 μM) (OctMc_P_). Subsequently, the NADH electron transfer-pathway state (FN) was studied by adding a high concentration of malate (2 mM, OctM_P_), pyruvate (5 mM, OctPM_P_), and glutamate (10 mM, OctPGM_P_). Then succinate (10 mM, OctPGMS_P_) was added to stimulate the S pathway (FNS), followed by glycerophosphate (10 mM, OctPGMSGp_P_) to reach convergent electron flow in the FNSGp-pathway to the Q-junction. Uncoupled respiration was next measured by performing a titration with CCCP (OctPGMSGp_E_), followed by inhibition of complex I (SGp_E_) with rotenone (0.5 μM, SGp_E_). Finally, residual oxygen consumption (ROX) was measured by adding Antimycin A (2.5 μM). ROX was then subtracted from all respiratory states, to obtain mitochondrial respiration. Results are expressed in pmol · s^−1^ · 1×10^6^ cells. The P/E control ratio, which reflects the control by coupling and limitation by the phosphorylation system, was subsequently calculated by dividing the OctPGMSGp_P_ value by the OctPGMSGp_E_ value.

### Radiolabelled glutamine uptake (protocol was adapted from^71^)

Two hiPSC lines (wibj_2 and oaqd_3) with 3 technical replicates each were compared to two hESC lines (SA121 and SA181) with 3 technical replicates of each. Both hiPSCs and hESCs were plated in 6-well plates 2 days before the transport assay (5×10^4^ cells/cm2 – this gives 1×10^6^ cells/well on “uptake day”). The cell growth media was carefully aspirated so as not to disturb the adherent monolayer of cells. They were washed gently by pipetting with 5 mls preheated (37°C) uptake solution (HBSS (pH 7.4), GIBCO) and aspirating off. This was repeated 3 times. They were then incubated with 0.5 ml of uptake solution containing [^3^H]glutamine (5 μCi/ml; Perkin Elmer, NET 55100) in either the presence, or absence, of L-glutamine (5 mM; Sigma) for 2 min.

Glutamine uptake was stopped by removing the uptake solution and washing cells with 2 ml of ice-cold stop solution (HBSS with 10 mM nonradioactive L-glutamine) three times. After the third wash, the cells were lysed in 200 μl of 0.1% SDS and 100 mM NaOH, and 100 μl was used to measure the radioactivity associated with the cells. Finally 100 μl sample was added to scintillation vials containing 3 mls scintillant (*OptiPhase HiSafe* 3, Perkin Elmer). β-radioactivity was measured with Tri-Carb 4910TR liquid scintillation counter.

The net glutamine CPM values were calculated by subtracting the Quench CPM values from the Glutamine CPM values.

### TEM Sample Preparation

wibj_2 and H1 cells were fixed on the dish in 4% paraformaldehyde and 2.5% glutaraldehyde in 0.1M sodium cacodylate buffer (pH 7.2) for 30 minutes then scraped and transferred to a tube and fixed for a further 30 minutes prior to pelleting. The pellets were cut into small pieces, washed 3 times in cacodylate buffer and then post-fixed in 1% OsO4 with 1.5% Na ferricyanide in cacodylate buffer for 60 min. After another 3 washes in cacodylate buffer they were contrasted with 1% tannic acid and 1% uranyl acetate. The cell pellets were then dehydrated through alcohol series into 100% ethanol, changed to propylene oxide left overnight in 50% propylene oxide 50% resin and finally embedded in 100% Durcupan resin (Sigma). The resin was polymerised at 60°C for 48hrs and sectioned on a Leica UCT ultramicrotome. Sections were contrasted with 3% aqueous uranyl acetate and Reynolds lead citrate before imaging on a JEOL 1200EX TEM using a SIS III camera.

### Proteomics search parameters

The data were searched and quantified with MaxQuant^72^ (version 1.6.7) against the human SwissProt database from UniProt^73^ (November 2019). The data were searched with the following parameters: type was set to Reporter ion on MS3 with 10plex TMT, stable modification of carbamidomethyl (C), variable modifications of oxidation (M), acetylation (proteins N terminus) and deamidation (NQ). The missed cleavage threshold was set to 2, and the minimum peptide length was set to 7 amino acids. The false discovery rate was set to 1% for positive identification at the protein and peptide spectrum match (PSM) level.

### Unique, shared and razor peptides

Peptides which are exclusive to a single protein group are considered unique peptides. Peptides whose sequences match more than one protein group are called shared peptides. Razor peptides are shared peptides whose intensity gets assigned to a single protein group despite matching multiple protein groups.

### Data filtering

All protein groups identified with less than either 2 razor or unique peptides or labelled as ‘Contaminant’, ‘Reverse’ or ‘Only identified by site’ were removed from the analysis.

### Peptide normalisation

For supplemental figures 2 & 3 peptide intensities were divided by the sum of the intensity from all histone peptides and were multiplied by 1×10^6^.

### Copy number calculations

Protein copy numbers were estimated following the “proteomic ruler” method^20^, but adapted to work with TMT MS3 data. The summed MS1 intensities were allocated to the different experimental conditions according to their fractional MS3 reporter intensities.

### Protein content estimations

The protein content was estimated using the following formula: CNL×LMW and then converting the data from Daltons to picograms, where CN is the protein copy number and MW is the protein molecular weight (in Da).

### Differential expression analysis

Fold changes and P-values were calculated in R. For individual proteins the p-values were calculated with the bioconductor package LIMMA^74^ version 3.7. The Q-values provided were generated in R using the “qvalue” package version 2.10.0. P-values for protein families and protein complexes were calculated in R using Welch’s T-test.

### hiPSC vs hESC overrepresentation analysis

All overrepresentation analysis were done on WebGestalt. The first analysis selected proteins with a fold change > 2 and a q-value < 0.001. The second analysis selected proteins whose fold change was lower than the median minus one standard deviation (0.195) and a q-value < 0.001. Both analyses used all identified proteins with 2 or more razor and unique peptides as a background and required an FDR lower than 0.05.

### Peptide coverage figures

The supplemental figures showing the peptide coverage across H1 and H2 histones (Fig. S2 & S3) were generated with Protter^75^.

## Data availability

The raw files and the mzTab outputs were uploaded to PRIDE as a full submission under the identifier PXD014502 and are available at https://www.ebi.ac.uk/pride/archive/projects/PXD014502

## Acknowledgements

We would like to thank Gabriel Sollberger well as all members of the Lamond Laboratory for their input and advice. This work was supported by the Wellcome Trust/MRC grant (098503/E/12/Z), Wellcome Trust grants (073980/Z/03/Z, 105024/Z/14/Z, 206293/Z/17/Z, 097418/Z/11/Z, 205023/Z/16/Z), BBSRC Project Grant (BB/V010948/1), EPSRC grant (EP/Y010655/1), a Wellcome Trust Equipment Award (202950/Z/16/Z) and a UK Research Partnership Infrastructure Fund award to the Centre for Translational and Interdisciplinary Research. Multiple figures in this paper were created using biorender.com

## Author contributions

A.J.B conceived the study, planned the experiments, analysed and interpreted the data. E.G executed all the proteomic sample preparation, and the mass spectrometry experiments. L.V.S performed the glutamine uptake assay. A.R.P performed the TEM experiments. F.S performed the respiration analysis. H.J. performed the EZQ assay and assisted with data interpretation. L.D cultured the hESC and hiPSCs, performed the pluripotency marker western blot and the cell cycle FACS experiments. C.E and E.K.J.H performed the histone western blots. H.Y., M.P and J.S helped to interpret the data. A.I.L, D.A.C and G.F supervised the project and helped to interpret the data. The paper written be A.J.B and A.I.L and edited by all authors.

## References

1 Smith, A. G. Embryo-derived stem cells: of mice and men. Annu Rev Cell Dev Biol 17, 435–462, doi:10.1146/annurev.cellbio.17.1.435 (2001).

2 Thomson, J. A. et al. Embryonic stem cell lines derived from human blastocysts. Science 282, 1145–1147, doi:10.1126/science.282.5391.1145 (1998).

3 Volarevic, V. et al. Ethical and Safety Issues of Stem Cell-Based Therapy. Int J Med Sci 15, 36–45, doi:10.7150/ijms.21666 (2018).

4 Takahashi, K. et al. Induction of pluripotent stem cells from adult human fibroblasts by defined factors. Cell 131, 861–872, doi:10.1016/j.cell.2007.11.019 (2007).

5 Takahashi, K. & Yamanaka, S. Induction of pluripotent stem cells from mouse embryonic and adult fibroblast cultures by defined factors. Cell 126, 663–676, doi:10.1016/j.cell.2006.07.024 (2006).

6 Kimbrel, E. A. & Lanza, R. Current status of pluripotent stem cells: moving the first therapies to the clinic. Nat Rev Drug Discov 14, 681–692, doi:10.1038/nrd4738 (2015).

7 Ebert, A. D. et al. Induced pluripotent stem cells from a spinal muscular atrophy patient. Nature 457, 277–280, doi:10.1038/nature07677 (2009).

8 Lee, G. et al. Modelling pathogenesis and treatment of familial dysautonomia using patient-specific iPSCs. Nature 461, 402–406, doi:10.1038/nature08320 (2009).

9 Liu, G. H. et al. Progressive degeneration of human neural stem cells caused by pathogenic LRRK2. Nature 491, 603–607, doi:10.1038/nature11557 (2012).

10 Mallon, B. S. et al. Comparison of the molecular profiles of human embryonic and induced pluripotent stem cells of isogenic origin. Stem Cell Res 12, 376–386, doi:10.1016/j.scr.2013.11.010 (2014).

11 Mallon, B. S. et al. StemCellDB: the human pluripotent stem cell database at the National Institutes of Health. Stem Cell Res 10, 57–66, doi:10.1016/j.scr.2012.09.002 (2013).

12 Guenther, M. G. et al. Chromatin structure and gene expression programs of human embryonic and induced pluripotent stem cells. Cell Stem Cell 7, 249–257, doi:10.1016/j.stem.2010.06.015 (2010).

13 Munoz, J. et al. The quantitative proteomes of human-induced pluripotent stem cells and embryonic stem cells. Mol Syst Biol 7, 550, doi:10.1038/msb.2011.84 (2011).

14 Phanstiel, D. H. et al. Proteomic and phosphoproteomic comparison of human ES and iPS cells. Nat Methods 8, 821–827, doi:10.1038/nmeth.1699 (2011).

15 Vitale, A. M. et al. Variability in the generation of induced pluripotent stem cells: importance for disease modeling. Stem Cells Transl Med 1, 641–650, doi:10.5966/sctm.2012-0043 (2012).

16 Kilpinen, H. et al. Common genetic variation drives molecular heterogeneity in human iPSCs. Nature 546, 370–375, doi:10.1038/nature22403 (2017).

17 Thompson, A. et al. Tandem mass tags: a novel quantification strategy for comparative analysis of complex protein mixtures by MS/MS. Anal Chem 75, 1895–1904 (2003).

18 McAlister, G. C. et al. MultiNotch MS3 enables accurate, sensitive, and multiplexed detection of differential expression across cancer cell line proteomes. Anal Chem 86, 7150–7158, doi:10.1021/ac502040v (2014).

19 Brenes, A., Hukelmann, J., Bensaddek, D. & Lamond, A. I. Multibatch TMT Reveals False Positives, Batch Effects and Missing Values. Mol Cell Proteomics 18, 1967–1980, doi:10.1074/mcp.RA119.001472 (2019).

20 Wisniewski, J. R., Hein, M. Y., Cox, J. & Mann, M. A “proteomic ruler” for protein copy number and concentration estimation without spike-in standards. Mol Cell Proteomics 13, 3497–3506, doi:10.1074/mcp.M113.037309 (2014).

21 Folmes, C. D. et al. Somatic oxidative bioenergetics transitions into pluripotency-dependent glycolysis to facilitate nuclear reprogramming. Cell Metab 14, 264–271, doi:10.1016/j.cmet.2011.06.011 (2011).

22 Howden, A. J. M. et al. Quantitative analysis of T cell proteomes and environmental sensors during T cell differentiation. Nat Immunol, doi:10.1038/s41590-019-0495-x (2019).

23 Marchingo, J. M., Sinclair, L. V., Howden, A. J. & Cantrell, D. A. Quantitative analysis of how Myc controls T cell proteomes and metabolic pathways during T cell activation. Elife 9, doi:10.7554/eLife.53725 (2020).

24 Turner, J. et al. Metabolic profiling and flux analysis of MEL-2 human embryonic stem cells during exponential growth at physiological and atmospheric oxygen concentrations. PLoS One 9, e112757, doi:10.1371/journal.pone.0112757 (2014).

25 Marchingo, J. M. & Cantrell, D. A. Protein synthesis, degradation, and energy metabolism in T cell immunity. Cell Mol Immunol 19, 303–315, doi:10.1038/s41423-021-00792-8 (2022).

26 Bhutia, Y. D. & Ganapathy, V. Glutamine transporters in mammalian cells and their functions in physiology and cancer. Biochim Biophys Acta 1863, 2531–2539, doi:10.1016/j.bbamcr.2015.12.017 (2016).

27 Broer, A., Rahimi, F. & Broer, S. Deletion of Amino Acid Transporter ASCT2 (SLC1A5) Reveals an Essential Role for Transporters SNAT1 (SLC38A1) and SNAT2 (SLC38A2) to Sustain Glutaminolysis in Cancer Cells. J Biol Chem 291, 13194–13205, doi:10.1074/jbc.M115.700534 (2016).

28 Marsboom, G. et al. Glutamine Metabolism Regulates the Pluripotency Transcription Factor OCT4. Cell Rep 16, 323–332, doi:10.1016/j.celrep.2016.05.089 (2016).

29 Tohyama, S. et al. Glutamine Oxidation Is Indispensable for Survival of Human Pluripotent Stem Cells. Cell Metab 23, 663–674, doi:10.1016/j.cmet.2016.03.001 (2016).

30 Eberle, D., Hegarty, B., Bossard, P., Ferre, P. & Foufelle, F. SREBP transcription factors: master regulators of lipid homeostasis. Biochimie 86, 839–848, doi:10.1016/j.biochi.2004.09.018 (2004).

31 Nose, F. et al. Crucial role of perilipin-3 (TIP47) in formation of lipid droplets and PGE2 production in HL-60-derived neutrophils. PLoS One 8, e71542, doi:10.1371/journal.pone.0071542 (2013).

32 Taanman, J. W. The mitochondrial genome: structure, transcription, translation and replication. Biochim Biophys Acta 1410, 103–123, doi:10.1016/s0005-2728(98)00161-3 (1999).

33 Nowinski, S. M. et al. Mitochondrial fatty acid synthesis coordinates oxidative metabolism in mammalian mitochondria. Elife 9, doi:10.7554/eLife.58041 (2020).

34 Mouw, J. K., Ou, G. & Weaver, V. M. Extracellular matrix assembly: a multiscale deconstruction. Nat Rev Mol Cell Biol 15, 771–785, doi:10.1038/nrm3902 (2014).

35 Romer, A. M. A., Thorseth, M. L. & Madsen, D. H. Immune Modulatory Properties of Collagen in Cancer. Front Immunol 12, 791453, doi:10.3389/fimmu.2021.791453 (2021).

36 Lanner, F. & Rossant, J. The role of FGF/Erk signaling in pluripotent cells. Development 137, 3351–3360, doi:10.1242/dev.050146 (2010).

37 Xu, R. H. et al. NANOG is a direct target of TGFbeta/activin-mediated SMAD signaling in human ESCs. Cell Stem Cell 3, 196–206, doi:10.1016/j.stem.2008.07.001 (2008).

38 Weinberger, L., Ayyash, M., Novershtern, N. & Hanna, J. H. Dynamic stem cell states: naive to primed pluripotency in rodents and humans. Nat Rev Mol Cell Biol 17, 155–169, doi:10.1038/nrm.2015.28 (2016).

39 Wang, X. et al. Reciprocal Signaling between Glioblastoma Stem Cells and Differentiated Tumor Cells Promotes Malignant Progression. Cell Stem Cell 22, 514–528 e515, doi:10.1016/j.stem.2018.03.011 (2018).

40 Filippou, P. S., Karagiannis, G. S. & Constantinidou, A. Midkine (MDK) growth factor: a key player in cancer progression and a promising therapeutic target. Oncogene 39, 2040–2054, doi:10.1038/s41388-019-1124-8 (2020).

41 Lu, Y. et al. Effect of midkine on gemcitabine resistance in biliary tract cancer. Int J Mol Med 41, 2003–2011, doi:10.3892/ijmm.2018.3399 (2018).

42 Vonwirth, V. et al. Inhibition of Arginase 1 Liberates Potent T Cell Immunostimulatory Activity of Human Neutrophil Granulocytes. Front Immunol 11, 617699, doi:10.3389/fimmu.2020.617699 (2020).

43 Wang, C. et al. CD276 expression enables squamous cell carcinoma stem cells to evade immune surveillance. Cell Stem Cell 28, 1597–1613 e1597, doi:10.1016/j.stem.2021.04.011 (2021).

44 Yue, G. et al. CD276 suppresses CAR-T cell function by promoting tumor cell glycolysis in esophageal squamous cell carcinoma. J Gastrointest Oncol 12, 38–51, doi:10.21037/jgo-21-50 (2021).

45 Pereira, B. I. et al. Senescent cells evade immune clearance via HLA-E-mediated NK and CD8(+) T cell inhibition. Nat Commun 10, 2387, doi:10.1038/s41467-019-10335-5 (2019).

46 Woodcock, C. L., Skoultchi, A. I. & Fan, Y. Role of linker histone in chromatin structure and function: H1 stoichiometry and nucleosome repeat length. Chromosome Res 14, 17–25, doi:10.1007/s10577-005-1024-3 (2006).

47 Robinson, P. J. & Rhodes, D. Structure of the ‘30 nm’ chromatin fibre: a key role for the linker histone. Curr Opin Struct Biol 16, 336–343, doi:10.1016/j.sbi.2006.05.007 (2006).

48 Cox, J. et al. Andromeda: a peptide search engine integrated into the MaxQuant environment. J Proteome Res 10, 1794–1805, doi:10.1021/pr101065j (2011).

49 Broer, S. Amino Acid Transporters as Targets for Cancer Therapy: Why, Where, When, and How. Int J Mol Sci 21, doi:10.3390/ijms21176156 (2020).

50 Ju, H. Q., Lin, J. F., Tian, T., Xie, D. & Xu, R. H. NADPH homeostasis in cancer: functions, mechanisms and therapeutic implications. Signal Transduct Target Ther 5, 231, doi:10.1038/s41392-020-00326-0 (2020).

51 Jin, L., Alesi, G. N. & Kang, S. Glutaminolysis as a target for cancer therapy. Oncogene 35, 3619–3625, doi:10.1038/onc.2015.447 (2016).

52 Wise, D. R. & Thompson, C. B. Glutamine addiction: a new therapeutic target in cancer. Trends Biochem Sci 35, 427–433, doi:10.1016/j.tibs.2010.05.003 (2010).

53 Reitzer, L. J., Wice, B. M. & Kennell, D. Evidence that glutamine, not sugar, is the major energy source for cultured HeLa cells. J Biol Chem 254, 2669–2676 (1979).

54 Brenes, A. J. et al. Erosion of human X chromosome inactivation causes major remodeling of the iPSC proteome. Cell Rep 35, 109032, doi:10.1016/j.celrep.2021.109032 (2021).

55 Lotz, S. et al. Sustained levels of FGF2 maintain undifferentiated stem cell cultures with biweekly feeding. PLoS One 8, e56289, doi:10.1371/journal.pone.0056289 (2013).

56 Ma, L., Chen, Z., Erdjument-Bromage, H., Tempst, P. & Pandolfi, P. P. Phosphorylation and functional inactivation of TSC2 by Erk implications for tuberous sclerosis and cancer pathogenesis. Cell 121, 179–193, doi:10.1016/j.cell.2005.02.031 (2005).

57 Pelletier, J., Graff, J., Ruggero, D. & Sonenberg, N. Targeting the eIF4F translation initiation complex: a critical nexus for cancer development. Cancer Res 75, 250–263, doi:10.1158/0008-5472.CAN-14-2789 (2015).

58 Galan, J. A. et al. Phosphoproteomic analysis identifies the tumor suppressor PDCD4 as a RSK substrate negatively regulated by 14-3-3. Proc Natl Acad Sci U S A 111, E2918–2927, doi:10.1073/pnas.1405601111 (2014).

59 Giulianelli, S. et al. FGF2 induces breast cancer growth through ligand-independent activation and recruitment of ERalpha and PRBDelta4 isoform to MYC regulatory sequences. Int J Cancer 145, 1874–1888, doi:10.1002/ijc.32252 (2019).

60 Li, Y., Guo, X. B., Wang, J. S., Wang, H. C. & Li, L. P. Function of fibroblast growth factor 2 in gastric cancer occurrence and prognosis. Mol Med Rep 21, 575–582, doi:10.3892/mmr.2019.10850 (2020).

61 Sooman, L. et al. FGF2 as a potential prognostic biomarker for proneural glioma patients. Acta Oncol 54, 385–394, doi:10.3109/0284186X.2014.951492 (2015).

62 Thomas, D. A. & Massague, J. TGF-beta directly targets cytotoxic T cell functions during tumor evasion of immune surveillance. Cancer Cell 8, 369–380, doi:10.1016/j.ccr.2005.10.012 (2005).

63 Li, M. O. & Flavell, R. A. TGF-beta: a master of all T cell trades. Cell 134, 392–404, doi:10.1016/j.cell.2008.07.025 (2008).

64 Munder, M. et al. Suppression of T-cell functions by human granulocyte arginase. Blood 108, 1627–1634, doi:10.1182/blood-2006-11-010389 (2006).

65 Lee, A. S., Tang, C., Rao, M. S., Weissman, I. L. & Wu, J. C. Tumorigenicity as a clinical hurdle for pluripotent stem cell therapies. Nat Med 19, 998–1004, doi:10.1038/nm.3267 (2013).

66 Muller, F. J. et al. A bioinformatic assay for pluripotency in human cells. Nat Methods 8, 315–317, doi:10.1038/nmeth.1580 (2011).

67 Ludwig, T. E. et al. Feeder-independent culture of human embryonic stem cells. Nat Methods 3, 637–646, doi:10.1038/nmeth902 (2006).

68 Hughes, C. S. et al. Ultrasensitive proteome analysis using paramagnetic bead technology. Mol Syst Biol 10, 757, doi:10.15252/msb.20145625 (2014).

69 Hughes, C. S. et al. Single-pot, solid-phase-enhanced sample preparation for proteomics experiments. Nat Protoc 14, 68–85, doi:10.1038/s41596-018-0082-x (2019).

70 Doerrier, C. et al. High-Resolution FluoRespirometry and OXPHOS Protocols for Human Cells, Permeabilized Fibers from Small Biopsies of Muscle, and Isolated Mitochondria. Methods Mol Biol 1782, 31–70, doi:10.1007/978-1-4939-7831-1_3 (2018).

71 Yeramian, A. et al. Arginine transport via cationic amino acid transporter 2 plays a critical regulatory role in classical or alternative activation of macrophages. J Immunol 176, 5918–5924, doi:10.4049/jimmunol.176.10.5918 (2006).

72 Cox, J. & Mann, M. MaxQuant enables high peptide identification rates, individualized p.p.b.-range mass accuracies and proteome-wide protein quantification. Nat Biotechnol 26, 1367–1372, doi:10.1038/nbt.1511 (2008).

73 The UniProt, C. UniProt: the universal protein knowledgebase. Nucleic Acids Res 45, D158–D169, doi:10.1093/nar/gkw1099 (2017).

74 Ritchie, M. E. et al. limma powers differential expression analyses for RNA-sequencing and microarray studies. Nucleic Acids Res 43, e47, doi:10.1093/nar/gkv007 (2015).

75 Omasits, U., Ahrens, C. H., Muller, S. & Wollscheid, B. Protter: interactive protein feature visualization and integration with experimental proteomic data. Bioinformatics 30, 884–886, doi:10.1093/bioinformatics/btt607 (2014).

