## Supplemental data for "Proteomic and functional comparison between human induced and embryonic stem cells"

[3] Present address: Centre for Inflammation Research, Institute for Regeneration and Repair, University of Edinburgh, Edinburgh, EH16 4UU, United Kingdom

[4] Present address: Drug Discovery Sciences, Boehringer Ingelheim Pharma GmbH & Co. KG, Biberach an der Riss, Germany

[8] Present address: Department of Physiology, Faculty of Medicine, Biomedical Center, University of Iceland, 101 Reykjavík, Iceland.

[9] Present address: Division of Cell Signalling, Fujii Memorial Institute of Medical Sciences, Institute of Advanced Medical Sciences, Tokushima University, 3-18-15 Kuramoto, Tokushima, 770-8503, Japan

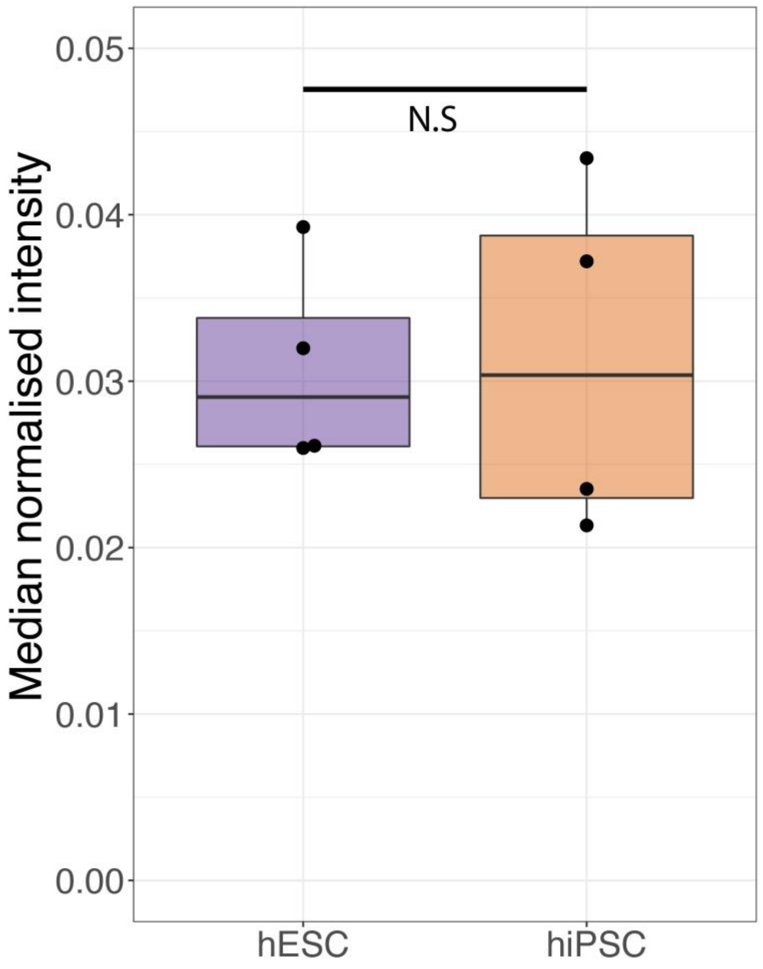


**Figure S1 – Histone intensity:** Box plot showing the sum of the normalised intensity for all histones across human embryonic stem cells (hESCs) and human induced pluripotent stem cells (hiPSCs). For the boxplots, the bottom and top hinges represent the 1st and 3rd quartiles. The top whisker extends from the hinge to the largest value no further than 1.5 3 IQR from the hinge; the bottom whisker extends from the hinge to the smallest value at most 1.5 3 IQR of the hinge.

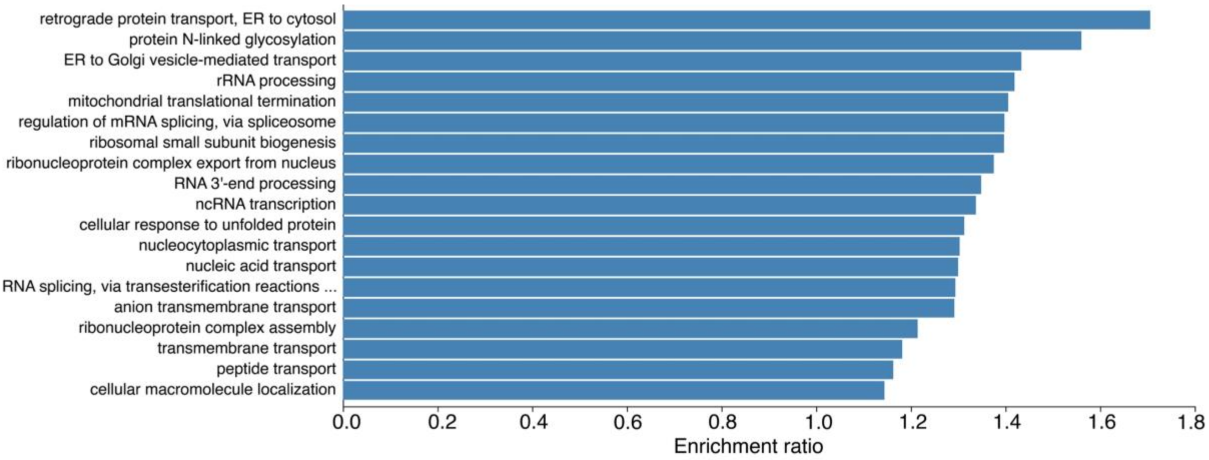


**Figure S2 – Proteins significantly increased in hiPSCs:** Bar plot showing the results of a Gene Ontology Biological Process overrepresentation analysis for proteins that are significantly increased in hiPSCs compared to hESCs.

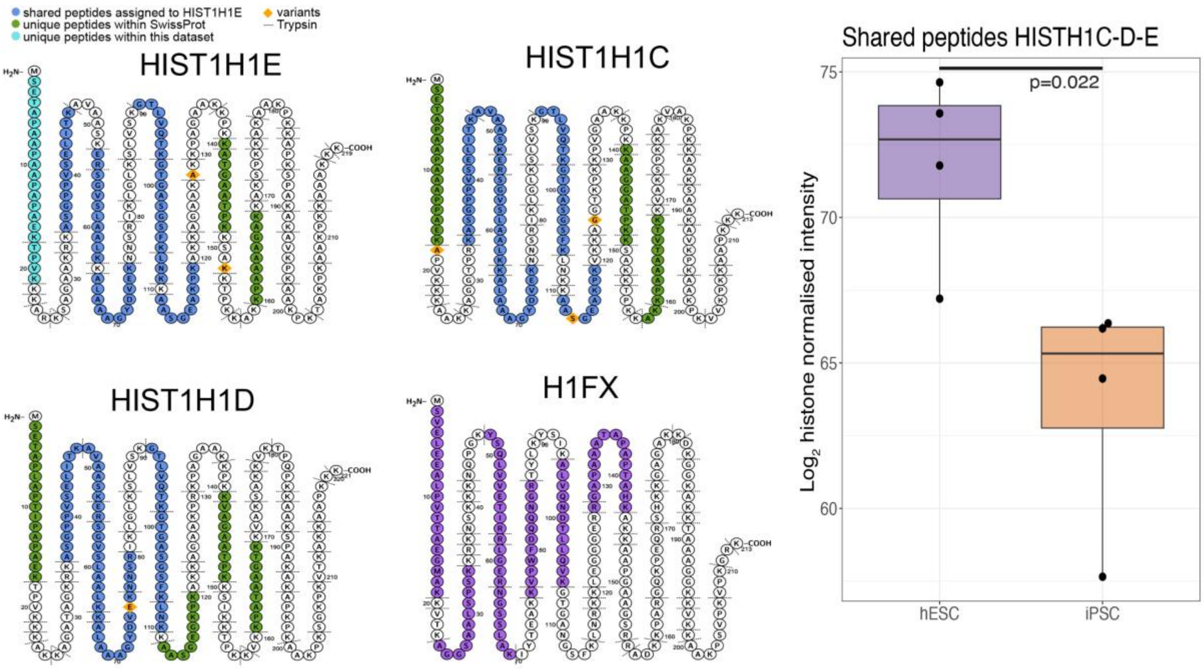


**Figure S3 – H1 histone peptides:** Schematic showing the unique and shared peptides detected for H1 histones as well as box plots showing the $Log_2$ histone normalised intensity for both hESC and hiPSCs. For all boxplots, the bottom and top hinges represent the 1st and 3rd quartiles. The top whisker extends from the hinge to the largest value no further than 1.5 3 IQR from the hinge; the bottom whisker extends from the hinge to the smallest value at most 1.5 3 IQR of the hinge.

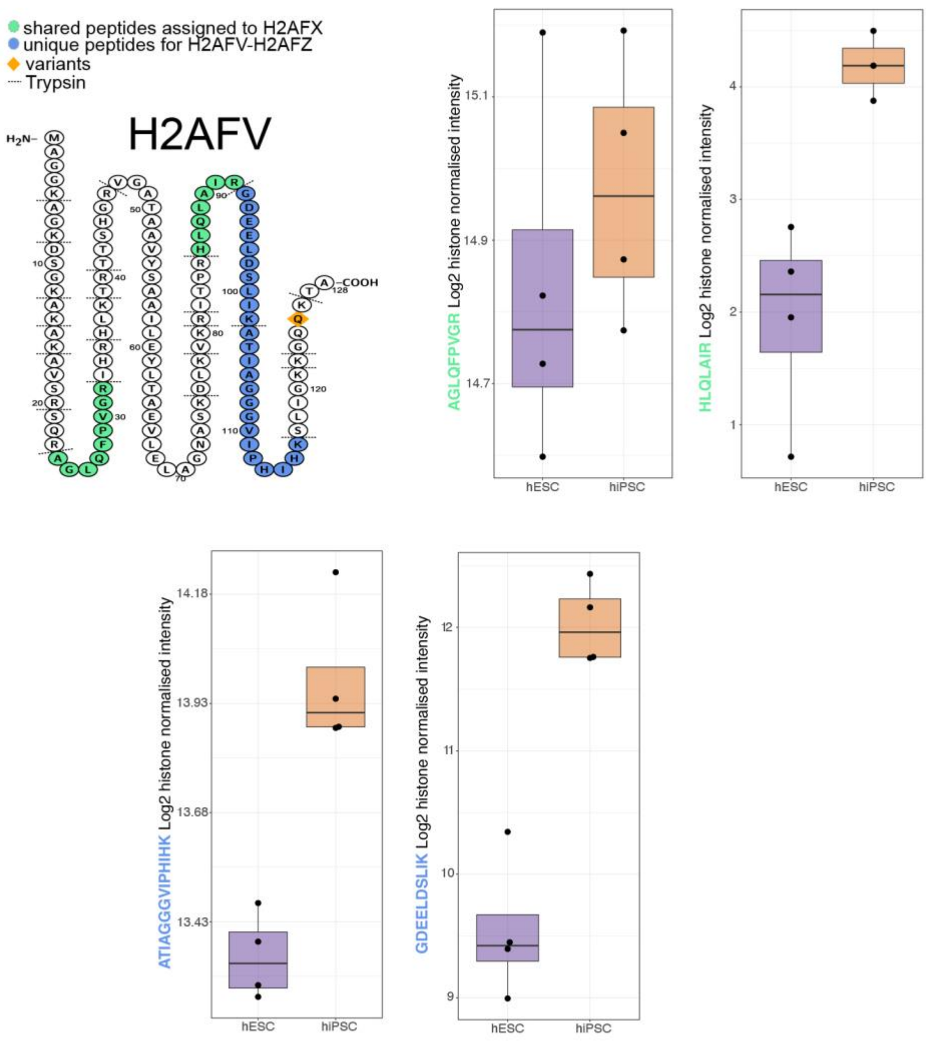


**Figure S4 – H2AFV peptides:** Schematic showing the unique and shared peptides detected for H2AFV as well as box plots showing the $Log_2$ histone normalised intensity for both hESC and hiPSCs. For all boxplots, the bottom and top hinges represent the 1st and 3rd quartiles. The top whisker extends from the hinge to the largest value no further than 1.5 3 IQR from the hinge; the bottom whisker extends from the hinge to the smallest value at most 1.5 3 IQR of the hinge.
